# PBN-PVT projection modulates negative affective states in mice

**DOI:** 10.1101/2021.03.11.434900

**Authors:** Ya-Bing Zhu, Yan Wang, Xiao-Xiao Hua, Ling Xu, Ming-Zhe Liu, Rui Zhang, Peng-Fei Liu, Jin-Bao Li, Ling Zhang, Di Mu

## Abstract

Long-lasting negative affections dampen enthusiasm for life, and dealing with negative affective states is essential for individual survival. The parabrachial nucleus (PBN) and the thalamic paraventricular nucleus (PVT) are critical for modulating affective states in mice. However, the functional role of the PBN-PVT projection in modulating affective states remains elusive. Here, we show that the PBN neurons send dense projection fibers to the PVT and form direct excitatory synapses with the PVT neurons. Activation of the PBN-PVT projection or PVT-projecting PBN neurons induces robust anxiety-like, aversion-like, and fear-like behaviors without affecting nociceptive behaviors. Inhibition of the PBN-PVT projection or PVT-projecting PBN neurons reduces fear-like and aversion-like behaviors. Furthermore, the PVT neurons innervated by the PBN are activated by aversive stimulation, and activation of PBN-PVT projection enhances the neuronal activity of PVT neurons in response to the aversive stimulus. Activation of these downstream PVT neurons induces anxiety-like behavior behaviors. Thus, our study indicates that the PBN-PVT projection modulates negative affective states in mice.

## Introduction

Threat and injury often induce defensive reactions, such as flight, freezing, hiding *(Öhman and Mineka, 2001)*, and negative affective states, such as fear and anxiety *(Jimenez et al., 2018)*. Such behavioral adaptations and psychological responses are essential for survival, and understanding the mechanisms is of fundamental interest. It is worth noting that the parabrachial nucleus (PBN) in the brainstem plays a critical role in encoding danger signals and promoting affective behavior states to limit harm in response to potential threats *(Campos et al., 2018)*.

The PBN receives the majority of the ascending inputs from the spinal cord *(Todd, 2010),* and the PBN neurons respond robustly to nociception, food neophobia, hypercapnia, and threat for maintaining homeostasis under stressful circumstances *(Campos et al., 2018; Kaur et al., 2013)*. The PBN relays this information (visceral malaise, taste, temperature, pain, itch) to brain areas, such as the hypothalamus, the central of the amygdala (CeA), thalamus, insular cortex (IC), and periaqueductal gray (PAG), to participate in diverse physiology process *(Chiang et al., 2019; Palmiter, 2018; Saper, 2016)*. A recent study has found that the subpopulations of PBN have distinct projection patterns and functions *(Chiang et al., 2020)*. The dorsal division PBN neurons projecting to the ventromedial hypothalamus (VMH) and PAG mediate escaping behaviors. In contrast, the external lateral division PBN neurons projecting to the bed nucleus of the stria terminalis (BNST) and the CeA mediate aversion and avoidance memory *(Chiang et al., 2020)*. Optogenetic manipulation of specific outputs from PBN generates a specific function *(Bowen et al., 2020)*. In the thalamus, the intralaminar thalamus nucleus (ILN) is the downstream target of PBN neurons that receive spinal cord inputs, and the ILN pathway participates in nociception processing *(Deng et al., 2020)*. Besides the ILN nucleus, the thalamic paraventricular nucleus (PVT) is another primary target of the PBN nucleus in the thalamus *(Chiang et al., 2020)*.

The PVT nucleus locates in the dorsal part of the midline thalamus *(Vertes et al., 2015)*, and has been heavily implicated in a range of affective behaviors *(Hsu et al., 2014)*. The functional roles of the PVT include diverse processes such as arousal (*Ren et al., 2018*), drug addiction (*Zhu et al., 2016*), reward-seeking (*Do-Monte et al., 2017*; *Engelke et al., 2021*), stress (*Beas et al., 2018*; *Gao et al., 2020*), associative learning and memory retrieval (*Penzo et al., 2015*; *Do-Monte et al., 2015*; *Zhu et al., 2018*; *Keyes et al., 2020*). The PVT receives a significant amount of inputs from the brainstem (locus coeruleus, PBN, PAG), hypothalamus, prefrontal cortical areas and projects to the infralimbic cortex, nucleus accumbens (NAc), BNST, and CeA *(Kirouac, 2015)*. The convergent signals included the arousal from the hypothalamus *(Ren et al., 2018)*, the emotional saliency from the prefrontal cortex *(Yamamuro et al., 2020)*, and the stress responsivity from locus coeruleus (LC) *(Beas et al., 2018)* might help to promote the appropriate behavioral responses to environmental challenges. However, despite the substantial improvements in our understanding of the PVT nucleus’s neurocircuitry, the functional role of the PBN-PVT projection remains mostly unknown.

In this study, we used virus tracing and electrophysiology to dissect the anatomical and functional connection between the PBN nucleus and the PVT nucleus. By using optogenetic and pharmacogenetic approaches, we then demonstrated that the PBN-PVT projection modulates negative affective states in mice.

## Results

### Functional connectivity pattern of the PBN-PVT projection

Previous studies have reported that the PVT could receive input from the PBN nucleus *(Chiang et al., 2020; Li and Kirouac, 2012)*. The detailed morphology of the PBN-PVT projection and whether these two nuclei form direct functional synapses remain unknown. We first injected the AAV2/8-hSyn-ChR2-mCherry virus into the PBN and found that there were dense projection fibers in the PVT (*Figure 1A−C*). We employed the whole-cell patch-clamp recording to examine the synaptic connectivity between the PBN and the PVT. Precisely time-locked action potentials were induced by a train of brief laser pulses (5 Hz, 10 Hz, and 20 Hz, *Figure 1G*). We found that optogenetic activation of the PBN projection fibers evoked excitatory postsynaptic currents (EPSCs) in 34 of 52 PVT neurons. The medial PVT showed higher connectivity (bregma -0.94 to -1.82 mm, 27 of 37 cells, 72.97%) than anterior PVT (bregma -0.22 to -0.94 mm, 2 of 6 cells, 33.33%) or posterior PVT (bregma -1.82 to -2.3 mm, 5 of 9 cells, 55.56%, *Figure 1D−F*). The average amplitude of the light-evoked EPSCs was 103.4 ± 11.93 pA (*Figure 1H*). Moreover, the latency of EPSCs was 3.13 ± 0.29 ms with a jitter of 0.16 ± 0.02 ms (*Figure 1I-J*), indicating monosynaptic connections between the PBN and the PVT nuclei. The EPSCs were sensitive to the Na^+^ channel blocker tetrodotoxin (TTX, 1 μM) and were rescued by the K^+^ channel blocker 4-aminopyridine (4-AP, 100 μM). The EPSCs were further blocked by the AMPA receptor antagonist NBQX (10 μM), confirming the monosynaptic glutamatergic innervation of the PVT neurons by the PBN neurons (*Figure 1K−L*). In addition, we also observed there were light-evoked inhibitory postsynaptic currents (IPSCs) in only 2 of 52 PVT neurons (less than 30 pA).

**Figure 1.**
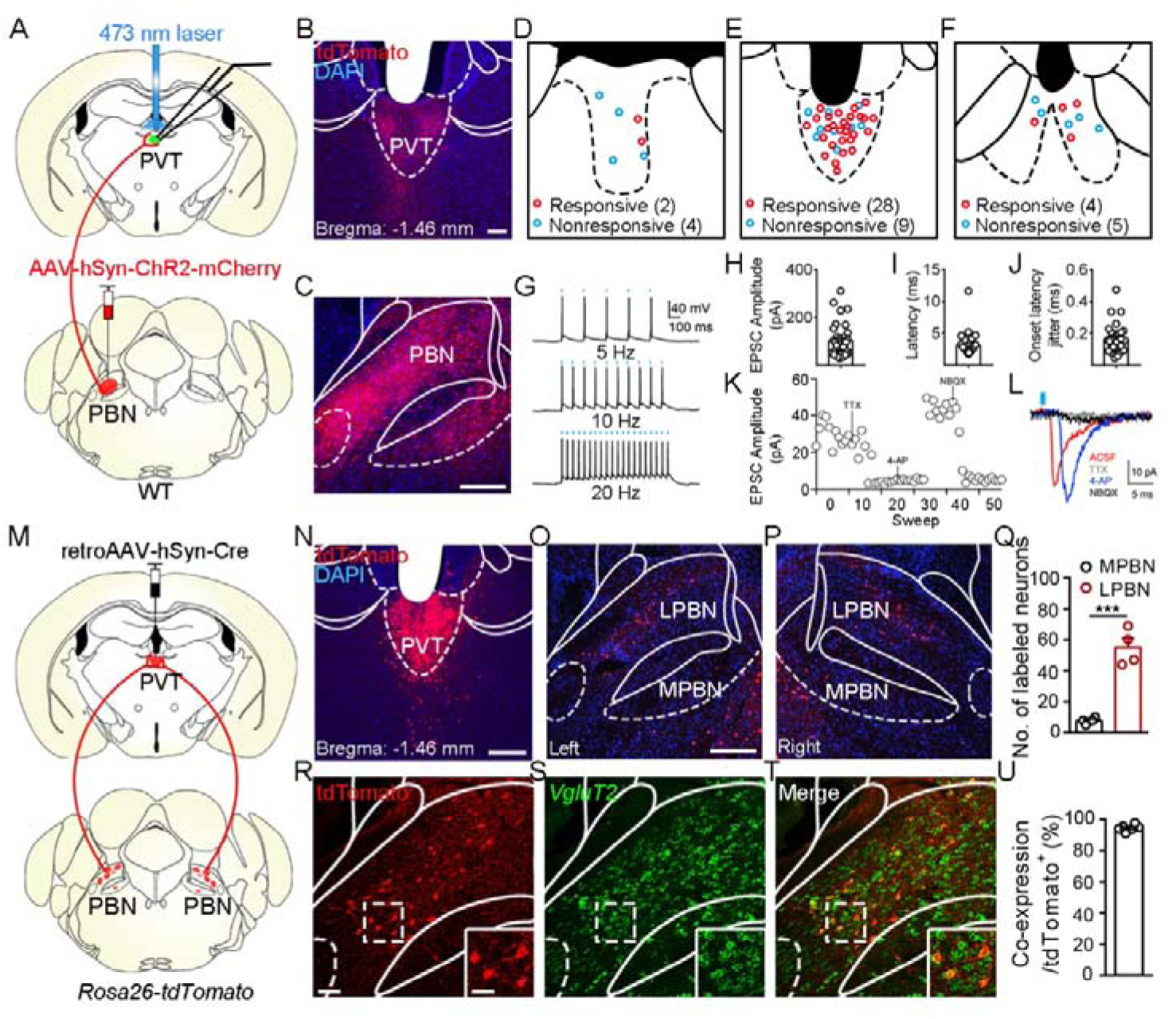
Functional connectivity pattern of the PBN-PVT projection. (A) The schematic for virus injection of AAV2/8-hSyn-ChR2-mCherry into the PBN nucleus and the slice recording with 473 nm laser stimulation. (B) The projection fibers in the PVT nucleus. Scale bar: 100 μm. (C) The AAV2/8-hSyn-ChR2-mCherry virus expression in the PBN nucleus. Scale bar: 200 μm. (D−F) The locations of the recorded cells in the anterior PVT (D), the middle PVT (E), and the posterior PVT (F). Red circles indicated neurons with excitatory postsynaptic currents (EPSCs), and blue circles indicated neurons without EPSCs. (G) The 473 nm laser-induced time-locked action potential firing at 5 Hz (top), 10 Hz (middle), and 20 Hz (bottom) in the ChR2-expressing neuron in the PBN. Scale bars: 100 ms, 40 mV. (H−J) The amplitude of light-evoked EPSCs (H), the latency of EPSCs (I), and the latency jitter of EPSCs (J) from all 34 responsive neurons in the PVT. (K) Amplitudes of light-evoked EPSCs recorded from a PVT neuron (right panel). (L) The light-evoked EPSC was completely blocked by 1 μM tetrodotoxin (TTX), rescued by 100 μM 4-aminopyridine (4-AP), and blocked by 10 μ (AMPA/kainate receptor antagonist). Scale bars: 5 ms, 10 pA. (M) Schematic shows retroAAV2/2-hSyn-Cre injection into the PVT nucleus on *Rosa26-tdTomato* mice. (N) The injection site in the PVT nucleus. Scale bar: 200 μm. (O and P) The distribution of the tdTomato positive neurons in the left PBN (O) and the right PBN (P). (Q) The quantification of the tdTomato positive neurons in the lateral PBN (LPBN) and the media PBN (MPBN). *n* = 4 mice. Scale bar: 200 μm. (R−T) Double staining of tdTomato with *VgluT2* mRNA by in situ hybridization. Scale bar: 50 μm, the scale bar in the quadrangle was 25 μm. (U) Quantification of the double-positive neurons over the total number of tdTomato positive neurons, *n* = 6 sections from 3 mice. ***p < 0.001, data were represented as mean ± SEM. Paired student’s *t*-test for Q. The following figure supplement is available for figure 1: **Figure 1−figure supplement 1.** Characterization of PVT-projecting neurons in the PBN nucleus. **Figure 1−figure supplement 2.** The distribution pattern of the PBN-PVT glutamatergic projection. **Figure 1−figure supplement 3.** The distribution pattern of collateral projection fibers from PVT-projecting PBN neurons.

Next, we injected the retroAAV2/2-hSyn-Cre virus into the PVT nucleus on *Rosa26-tdTomato* mice which could retrogradely label projection neurons in the PBN (*Figure 1M−P*). We found that tdTomato^+^ neurons were bilaterally located in the lateral PBN (55 ± 5.82 neurons, *n* = 4 mice) and rarely in the medial PBN (7.75 ± 1.03 neurons, *Figure 1O−Q*). These results indicate that bilateral PBN project to the PVT. We then performed tdTomato staining with *type 2 vesicular glutamate transporter* (*VgluT2)* mRNA in situ hybridization and found that about 94.4% of tdTomato neurons express *VgluT2 mRNA* (*Figure 1R-U*). These results indicate that the majority of PVT-projecting PBN neurons are glutamatergic. We also examined several markers for subpopulations of PBN neurons, including tachykinin 1 receptor (Tacr1), tachykinin 1 (Tac1), prodynorphin (Pdyn), calcitonin gene-related peptide (CGRP). And we found that tdTomato neurons were only partially co-labeled with *Tacr1*, *Tac1,* or *Pdyn* mRNA, but not with CGRP (*Figure 1−figure supplement 1*).

Then, we used the *VgluT2-ires-Cre* mice combined with the AAV2/8-EF1a-DIO-EGFP virus to specifically label the glutamatergic neurons of the PBN nucleus (*Figure 1−figure supplement 2A*). Robust expression of AAV2/8-EF1a-DIO-EGFP was found in both the lateral and medial PBN nuclei (*Figure1−figure supplement 2B−D*). It is worth noting that the density of EGFP^+^ fibers was higher in the middle and posterior PVT (*Figure1−figure supplement 2E−H*), considering that the notion of the posterior PVT being a particularly aversive region of the PVT (*Gao et al., 2020*). We also examined the collateral projections from PVT-projecting PBN neurons (*Figure 1−figure supplement 3*). The collateral projections were also found in the BNST, lateral hypothalamus (LH), paraventricular nucleus of the hypothalamus (PVN), PAG, but not CeA or VMH.

### Optogenetic activation of PBN-PVT projection induces anxiety-like behaviors and aversion-like behaviors

We injected the AAV2/9-EF1a-DIO-ChR2-mCherry virus or AAV2/9-EF1a-DIO-mCherry virus into the bilateral PBN of *VgluT2-ires-Cre* mice and implanted optic fibers above the PVT to activate the PBN-PVT projection selectively (*Figure 2A*). Four weeks after surgery, we found robust expression of ChR2-mCherry (*Figure 2B−C, Figure 2−figure supplement 1A*) or mCherry (*Figure 2−figure supplement 1C*) in bilateral PBN neurons and the axon terminals in the PVT (*Figure 2D, Figure 2−figure supplement 1B, Figure 2−figure supplement 1D*).

**Figure 2.**
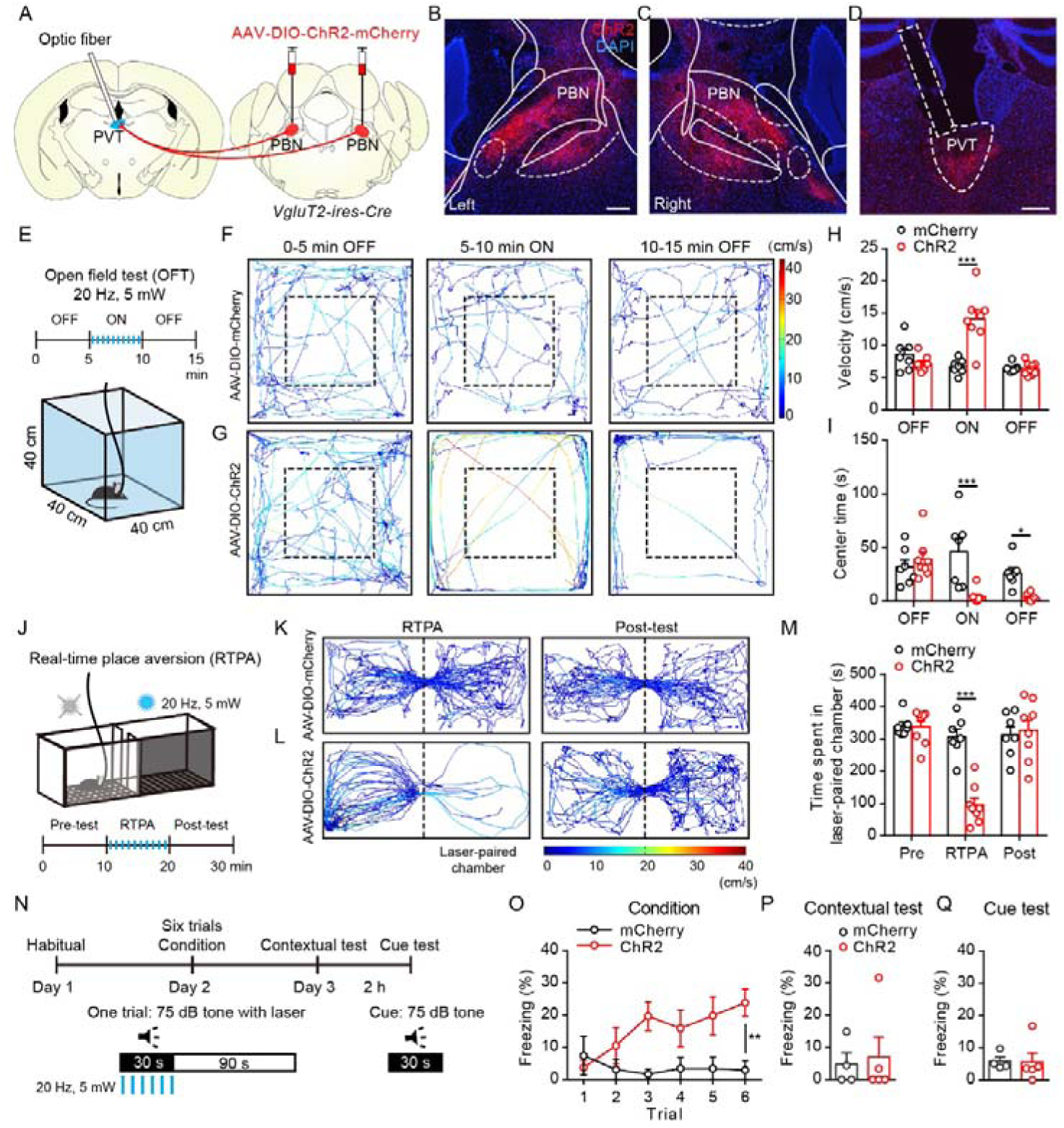
Optogenetic activation of the PBN-PVT projection induced negative affective states. (A) The illustration showed the injection of the AAV2/9-EF1a-DIO-ChR2-mCherry virus into the PBN nucleus and the optic fiber above the PVT on the *VgluT2-ires-Cre* mice. (B and C) The virus injection sites of the left PBN (B) and the right PBN (C). Scale bar: 200 μm. (D) The projection axons from the PBN and the location of the optic fiber (rectangle) in the PVT. Scale bar: 200 μm. (E) The schematic of the open field test (OFT) with optogenetic activation via a 473 nm laser. (F and G) The example traces of the 15 minutes optogenetic manipulation OFT from an AAV2/9-EF1a-DIO-mCherry virus injected mouse (F) or an AAV2/9-EF1a-DIO-ChR2-mCherry virus injected mouse (G). (H and I) Quantification of the velocity (H) and the center time (I) in the OFT, mCherry group: *n* = 7 mice; ChR2 group: *n* = 8 mice. (J) The illustration of the real-time place aversion test (RTPA) with optogenetic activation via a 473 nm laser. The right side was paired with the laser. (K and L) The example traces of the RTPA and post-test from the mice injected with AAV2/9-EF1a-DIO-mCherry (K) or AAV2/9-EF1a-DIO-ChR2-mCherry (L). (M) Quantification of the time spent in the laser-paired chamber in the pre-test (Pre), RTPA, and post-test (Post), mCherry group: *n* = 7 mice; ChR2 group: *n* = 8 mice. (N) Schematic timeline of cue-dependent optogenetic conditioning. (O) Conditioned-freezing responses to sound cue paired with optogenetic activation of the PBN-PVT projection during training, mCherry group: *n* = 4 mice; ChR2 group: *n* = 5 mice. (P and Q) Optogenetic activation of the projection fibers from the PBN in the PVT did not induce context-dependent fear (P) and cue-dependent fear (Q), mCherry group: *n* = 4 mice; ChR2 group: *n* = 5 mice. *p < 0.05, **p < 0.01, ***p < 0.001, all data were represented as mean ± SEM. Two-way ANOVA followed by Bonferroni test for H, I, M, and O. Unpaired student’s *t*-test for P and Q. The following figure supplement is available for figure 2: **Figure 2−figure supplement 1.** The virus expression in the PBN and the optic fiber position in the PVT of *VgluT2-ires-Cre* mice injected with AAV2/9-EF1a-DIO-ChR2-mCherry virus or AAV2/9-EF1a-DIO-mCherry virus. **Figure 2−figure supplement 2.** Effects of optogenetic activation of PBN-PVT projection in the OFT and the CPA. **Figure 2−video 1.** Optogenetic activation of PBN-PVT projection in OFT. The 473 nm laser (20 Hz, 5 ms, 5 mW) was delivered from 00:10 to 05:10 in the video. **Figure 2−video 2.** Optogenetic activation of PBN-PVT projection in RTPA. The 473 nm laser (20 Hz, 5 ms, 5 mW) was delivered when the mouse entered the laser-paired chamber and withdrew when the mouse exited the laser-paired chamber during the 10 minutes. The video was played with 4x speed.

We performed a 15 minutes optogenetic manipulation open field test (OFT, 0−5 minutes laser OFF, 5−10 minutes laser ON, 10−15 minutes laser OFF, *Figure 2E*). Optogenetic activation (473 nm, 20 Hz, 5 mW, 5ms) of the efferents from the PBN to the PVT elicited instant running behavior along the chamber wall with a significantly increased velocity in ChR2 group mice (*Figure 2F−H, Figure 2−video 1*). The ChR2 injected mice rarely entered into the center area of the chamber, represented as a decrease in center time than that of the control group (*Figure 2I*). It is worth noting that the velocity returned to normal once the laser was off, but the time spent in the center was still lower than the control group in the 5 minutes after stimulation. These results indicated that the anxiety could last for at least several minutes after acute activation. Although the speed increased during the laser ON period, the unmoving time of the ChR2 mice during the laser ON period was also increased (*Figure 2−figure supplement 2A*). Therefore, the distance during the laser ON period and the total distance in 15 minutes were not changed (*Figure 2−figure supplement 2B*).

To dissect a more detailed profile of the behaviors in the OFT, we further divided the laser ON period (5−10 minutes) into 5 one-minute periods and analyzed the velocity, unmoving time, center time, distance, and jumping (*Figure 2−figure supplement 2C−G*). We found that the velocity and unmoving time were increased, and the center time was decreased in the ChR2 mice during most periods (*Figure 2−figure supplement 2C−E*). Furthermore, we observed that the distance and jumping behaviors were increased mainly in the first one-minute period in ChR2 mice (*Figure 2−figure supplement 2F−G*). This detailed analysis indicates that optogenetic activation induces brief and robust running, jumping behaviors, and persistent anxiety-like behaviors, such as unmoving and less time spent in the center.

Besides anxiety, another critical component of negative affective states is aversion. Therefore, we used the real-time placed aversion test (RTPA) to explore the function of optogenetic activation of the PBN-PVT projection (*Figure 2J*). We found that the ChR2 mice spent less time in the laser-paired chamber, and the aversion disappeared when the laser was off (*Figure 2K−M, Figure 2−video 2*). We also used a prolonged conditioning protocol that mimics drug-induced conditioned place aversion (*Figure 2−figure supplement 2H*). We found that the ChR2 mice did not display aversion in the post-conditioning test (*Figure 2-figure supplement 2I*). These results indicate that activation of the PBN-PVT projection is sufficient to induce aversion but could not enable associative aversive memory formation.

To further confirm this instant aversion phenomenon, we subjected mice to the cue-dependent optogenetic conditioning test (*Figure 2N*). A 30 seconds auditory conditioning stimulus (CS) co-terminated with 30 seconds of synchronous optogenetic activation of the PBN-PVT projection (laser stimulus, LS) in this test. The ChR2 expressing mice generated significant freezing behavior during 6 CS-LS pairings (*Figure 2O*). However, the freezing behavior to the same context or to the auditory cue in a novel context was disappeared on the second day (*Figure 2P−Q*). These results demonstrate that optogenetic activation of the PBN-PVT projection induces instant aversion and freezing but does not drive associative fear memory formation.

### Pharmacogenetic activation of the PVT-projecting PBN neurons induces anxiety-like behaviors and freezing behaviors

We also used pharmacogenetic manipulation and retrograde tracing to confirm the effects of activating the PVT-projecting PBN neurons. We first injected the retroAAV2/2-hSyn-Cre virus into the PVT, and the AAV2/9-hSyn-DIO-hM3Dq-mCherry virus or control virus into bilateral PBN to specifically transduce the PVT-projecting PBN neurons with a designer receptor exclusively activated by designer drugs (DREADDs, *Figure 3A*) *(Armbruster et al., 2007)*. The PVT-projecting PBN neurons could be activated by intraperitoneal injection of clozapine N-oxide dihydrochloride (CNO, *Figure 3B−D*). The region of the virus expression in the PBN nucleus is shown in *Figure 3−figure supplement 1*. Consistent with the optogenetic activation results, the hM3Dq mice spent less time in the center, had more unmoving time, and traveled fewer distances than the mCherry mice in the OFT (*Figure 3E−I*). At the same time, the velocities were not significantly different (*Figure 3J*). We also found that activation of the PVT-projecting PBN neurons did not affect motor ability in the rotarod test (*Figure 3−figure supplement 2G*). Besides, the hM3Dq mice showed decreased exploration time of open quadrants in the EZM (*Figure 3K−L*), further suggesting that activation of PVT-projecting PBN neurons could induce anxiety-like behaviors.

**Figure 3.**
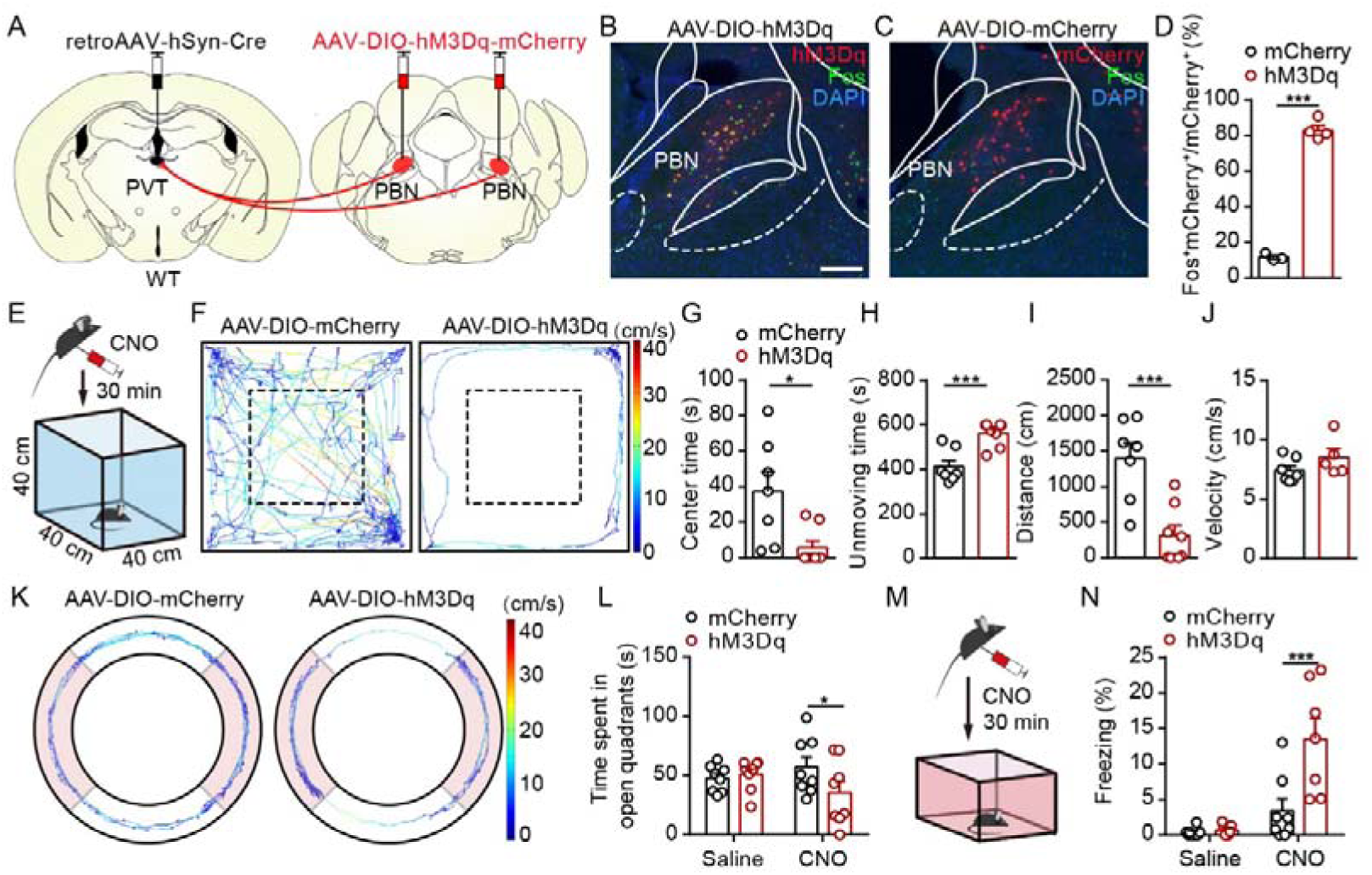
Pharmacogenetic activation of the PVT-projecting PBN neurons induced anxiety-like behaviors and fear-like behaviors. (A) The illustration showed virus injection of retroAAV2/2-hSyn-Cre into the PVT nucleus and bilateral injection of AAV2/9-hSyn-DIO-hM3Dq-mCherry into the PBN nucleus. (B and C) CNO administration evokes Fos expression in AAV2/9-hSyn-DIO-hM3Dq-mCherry injected mice (B) but not in AAV2/9-EF1a-DIO-mCherry injected mice (C). Scale bar: 200 μm. (D) Percentage of co-labeled neurons in the PBN, mCherry group: *n* = 3 mice; hM3Dq group: *n* = 4 mice. (E) The illustration of the OFT test with pharmacogenetic activation. (F) Example of the OFT traces from the mice infected with AAV2/9-EF1a-DIO-mCherry or AAV2/9-hSyn-DIO-hM3Dq-mCherry. (G−I) Quantification of the center time (G), the unmoving time (H), the total distance (I) in the OFT, mCherry group: *n* = 7 mice; hM3Dq group: *n* = 8 mice. (J) Quantification of the velocity in the OFT, mCherry group: *n* = 7 mice; hM3Dq group: *n* = 5 mice. (K) Example elevated zero maze (EZM) traces from the mice infected with AAV2/9-EF1a-DIO-mCherry and AAV2/9-hSyn-DIO-hM3Dq-mCherry. (L) Quantification of the time spent in open quadrants in the EZM test, *n* = 8 mice per group. (M) The illustration of pharmacogenetic activation-induced fear-like freezing behavior. (N) Pharmacogenetic activation of PVT-projecting PBN neurons induced fear-like freezing behaviors, mCherry group: *n* = 8 mice; hM3Dq group: *n* = 7 mice. *p < 0.05, ***p < 0.001, all data were represented as mean ± SEM. Unpaired student’s *t*-test for D, G, H, I, and J. Two-way ANOVA followed by Bonferroni test for L and N. The following figure supplements are available for figure 3: **Figure 3−figure supplement 1.** The virus expression in the PBN of mice injected with AAV2/9-hSyn-DIO-hM3Dq-mCherry or AAV2/9-EF1a-DIO-mCherry in the pharmacogenetic manipulation. **Figure 3−figure supplement 2.** Pharmacogenetic activation of PVT-projecting PBN neurons did not affect depressive-like behaviors, basal nociceptive thresholds, formalin-induced licking behavior, or motor function.

We further evaluated freezing behaviors in the fear conditioning chamber and found that the hM3Dq mice displayed more freezing behaviors after injection of CNO than control mice (*Figure 3M−N*). Although activation of PVT-projecting PBN neurons induced significant anxiety-like behavior, it did not affect the depressive-like behaviors evaluated by the tail suspension test (TST, *Figure 3−figure supplement 2A*) and the forced swimming test (FST, *Figure 3−figure supplement 2B*). Previous studies have revealed that the PBN receives direct projections from the spinal cord and plays a vital role in pain processing *(Deng et al., 2020; Sun et al., 2020)*. We then assessed whether pharmacogenetic activation of the PVT-projecting PBN neurons affects the nociceptive behaviors. By performing the von Frey test and Hargreaves test, we found that the basal nociceptive thresholds were not affected after pharmacogenetic activation of PVT-projecting PBN neurons (*Figure 3−figure supplement 2C−D*). Given that the distinct mechanisms between the reflexive and coping responses induced by nociceptive stimulation *(Huang et al., 2019)*, we injected formalin into the paw to induce inflammatory pain. We found that activation of the PVT-projecting PBN neurons did not affect the formalin-evoked licking behaviors (*Figure 3−figure supplement 2E−F*). These results indicate that the PBN-PVT projection might not participate in the pain signal processing.

### Inhibition of the PBN-PVT projection reduces the 2-MT-induced aversive behaviors and footshock-induced fear behaviors

The activation manipulation results prompted us to investigate whether inhibition of the PBN-PVT projection could modulate the negative affective states. We first injected the AAV2/9-EF1a-DIO-NpHR3.0-EYFP virus or the AAV2/8-EF1a-DIO-EGFP virus into the PBN and implanted the optic fibers into the PVT of *VgluT2-ires-Cre* mice (*Figure 4A−C, Figure 4−figure supplement 1*). We used 2-methyl-2-thiazoline (2-MT), a widely-used odorant molecule that could generate innate fear-like freezing responses in rodents *(Isosaka et al., 2015)*, to induce the fear-like state. We found that 589 nm laser-induced inhibition of the PBN-PVT projection reduced the aversion caused by the 2-MT (*Figure 4D−F*) and increased the moving duration (*Figure 4G*). We also observed that inhibition of the PBN-PVT projection increased the time in the 2-MT zone in the OFT (*Figure 4H−I*). Besides the 2-MT, footshock is another paradigm that induces robust freezing behaviors. We found that constant inhibition of the PBN-PVT projection reduced the footshock-induced freezing behaviors (*Figure 4J−K*).

**Figure 4.**
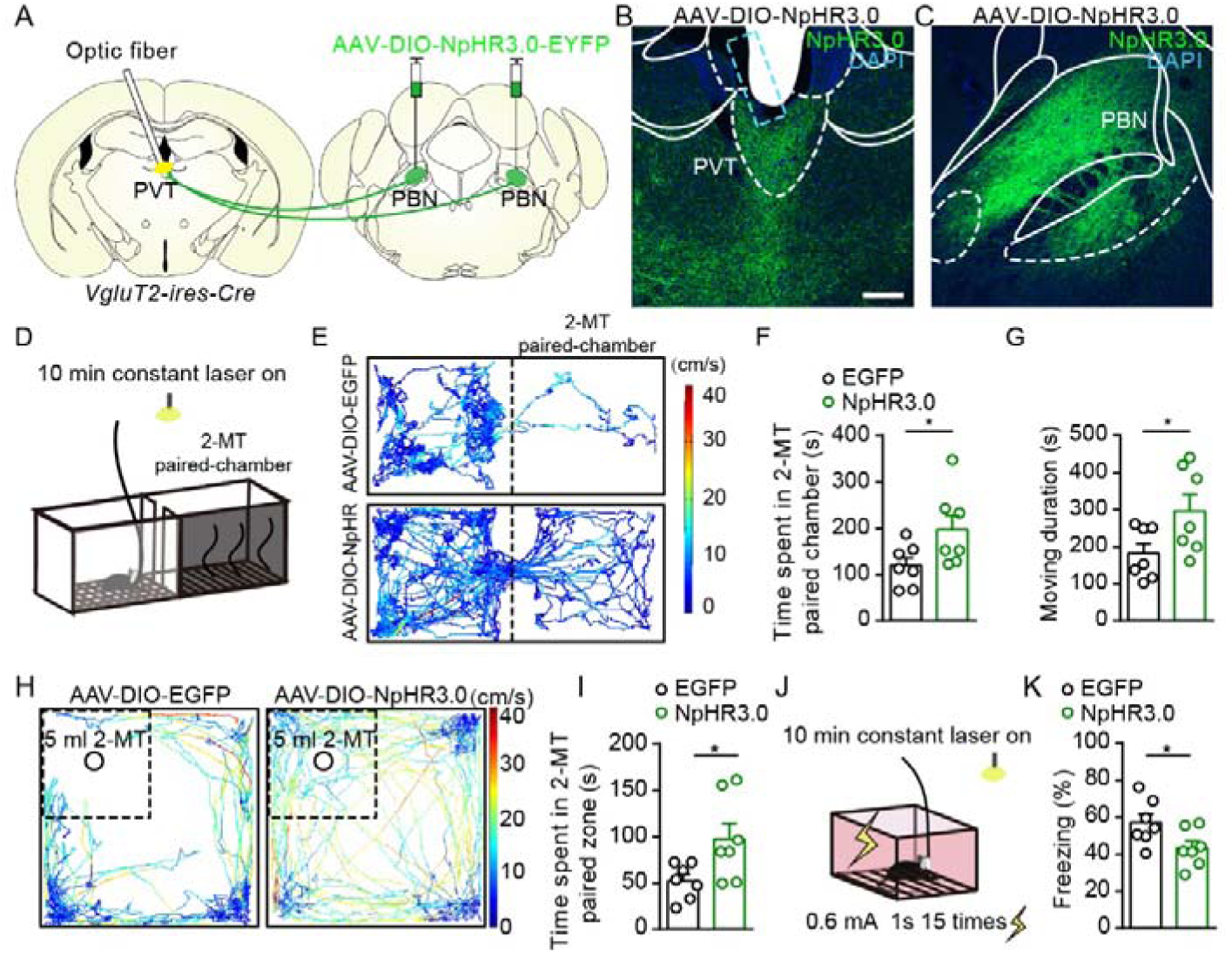
Optogenetic inhibition of the PBN-PVT projection reduced aversion-like behavior and fear-like behaviors. (A) The illustration showed the bilateral injection of AAV2/9-EF1a-DIO-NpHR3.0-EYFP virus into the PBN and placement of optic fiber above the PVT on *VgluT2-ires-Cre* mice. (B and C) Examples of AAV2/9-EF1a-DIO-NpHR3.0-EYFP expression in the PVT (B) and PBN (C). The cyan rectangle represented the position of the optic fiber. Scale bar: 200 μm. (D) Schematic of 2-MT induced aversion test with optogenetic inhibition via the 589 nm laser. (E) Representative traces of the mice infected with AAV2/8-EF1a-DIO-EGFP or AAV2/9-EF1a-DIO-NpHR3.0-EYFP in two chambers. (F and G) Quantification of the time spent in the 2-MT paired chamber (F) and the total moving duration (G), *n* = 7 mice per group. *p < 0.05, **p < 0.01, (H) Representative traces of the mice infected with AAV2/8-EF1a-DIO-EGFP or AAV2/9-EF1a-DIO-NpHR3.0-EYFP in the OFT chamber. (I) Quantification of the time spent in the 2-MT zone, *n* = 7 mice per group. (J) Illustration of footshock-induced freezing behavior with optogenetic inhibition via a 589 nm laser. (K) Quantification of the freezing behavior, *n* = 7 mice per group. *p < 0.05, all data were represented as mean ± SEM. Unpaired student’s *t*-test for F, G, I, and K. The following figure supplements are available for figure 4: **Figure 4−figure supplement 1.** The virus expression in the PBN and the optic fiber position in the PVT of *VgluT2-ires-Cre* mice injected with AAV2/9-EF1a-DIO-NpHR3.0-EYFP or AAV2/8-EF1a-DIO-EGFP. **Figure 4−figure supplement 2.** Optogenetic inhibition of the PBN-PVT projection did not affect associative fear memory acquisition and retrieval. **Figure 4−figure supplement 3.** Optogenetic inhibition of the PVT-projecting PBN neurons reduced the aversion-like behavior and fear-like freezing behavior.

We also examined whether inhibition of the PBN-PVT projection affects aversive memory acquisition or retrieval (*Figure 4−figure supplement 2A*). We briefly suppressed the activity of the PBN-PVT projection during footshock stimulation and found that freezing levels were not changed (*Figure 4−figure supplement 2B*). We further compared the freezing levels in contextual and cue tests without or with laser and found that aversive memory retrieval was not affected either (*Figure 4−figure supplement 2C−D*). In addition, we performed optogenetic inhibition of the PVT-projecting PBN neurons and observed similar phenomena (*Figure 4−figure supplement 3*).

### PBN input shapes PVT neuronal responses to aversive stimulation

By using *in vivo* fiber photometry, we found that the calcium signals of the PVT neurons were increased after aversive stimuli, such as footshock and air puff (*Figure 5−figure supplement 1*), indicating that the calcium signal of the PVT neurons is increased in response to aversive stimuli. Besides, we injected the AAV2/1-hSyn-Cre virus, which could anterogradely label the downstream neurons *(Zingg et al., 2017)*, into the PBN of *Rosa26-tdTomato* mice (*Figure 5A*). The distribution pattern of tdTomato^+^ neurons in PVT (hereafter referred to as PVT_PBN_ neurons) was shown in *Figure 5B−D*. We used Fos as a marker to assess the activity change in 2-MT treated mice and footshock treated mice. The percentage of Fos^+^tdTomato^+^ neurons/tdTomato^+^ neurons in the PVT was significantly increased in the aversive stimuli-treated mice than that of control mice (*Figure 5E−H*), confirming that the PVT_PBN_ neurons could be activated by aversive stimuli.

**Figure 5.**
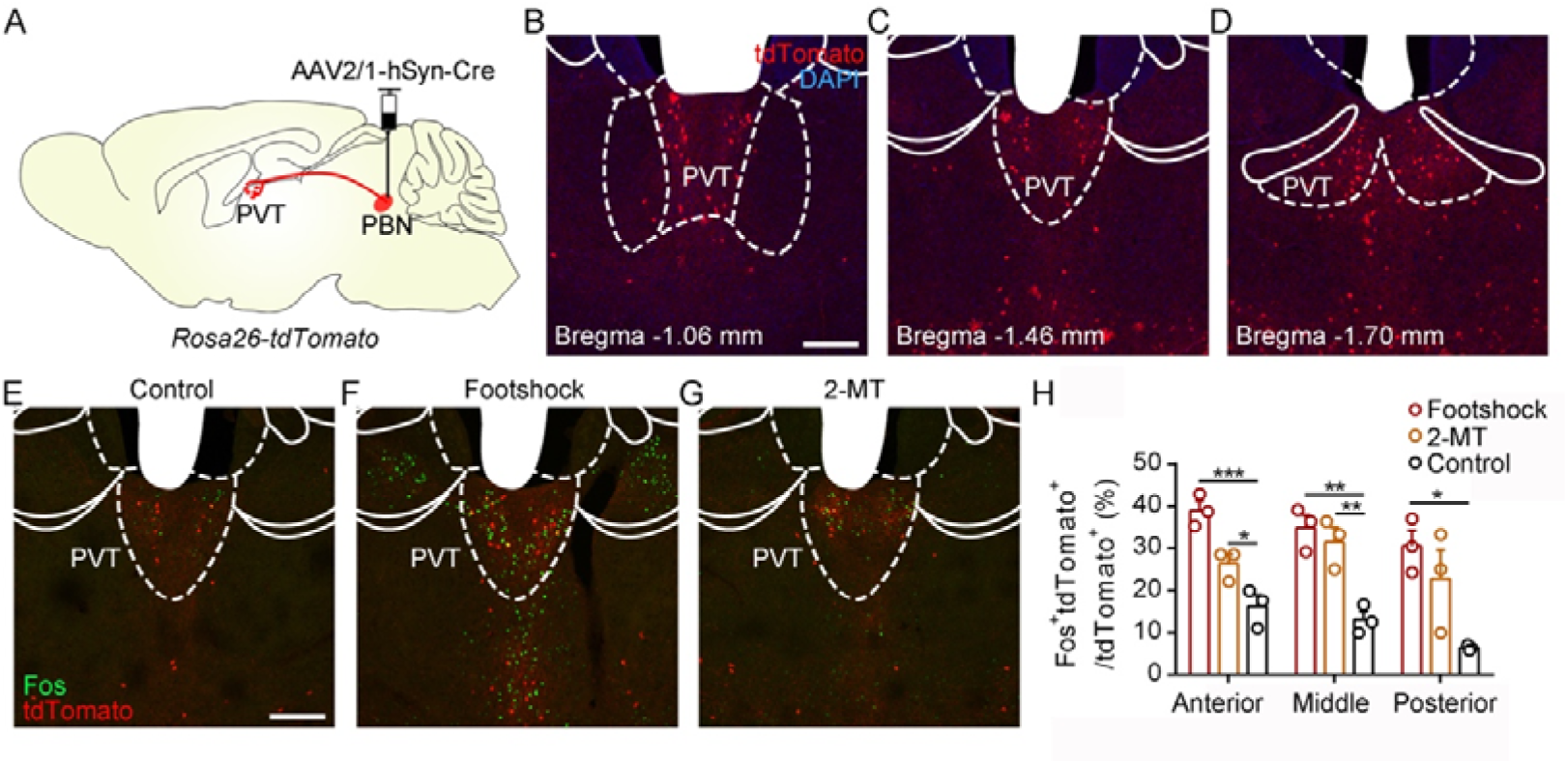
Activation of PVT_PBN_ by diverse aversive stimuli. (A) The illustration showed the injection of AAV2/1-hSyn-Cre into the PBN of *Rosa26-tdTomato* mice. (B−D) The distribution of the neurons in the PVT at bregma -1.06 mm(B), bregma -1.46 mm (C), and bregma -1.70 mm (D). Scale bar: 200 μm. (E−G) Fos induced by habituation control (E), footshock (F), or 2-MT (G) co-labeled with the tdTomato positive neurons in the PVT. Scale bar: 200 μm. (H) Quantification of the co-labeled neurons, *n* = 3 mice per group. *p < 0.05, **p < 0.01, ***p < 0.001, all data were represented as mean ± SEM, one-way ANOVA followed by Bonferroni test for H. The following figure supplements are available for figure 5: **Figure 5−figure supplement 1.** Calcium signals of PVT neurons in response to aversive stimuli.

The next question is whether the PBN-PVT projection modulates the neuronal activity of the PVT neurons in response to aversive stimuli. We first injected the AAV2/9-EF1a-DIO-ChR2-mCherry virus into the PBN and performed a dual Fos staining (*Seike et al.,* 2020), detecting *fos* mRNA and Fos protein induced by two episodes of stimulation (*Figure 6−figure supplement 1A*). We found that there was a broad overlap between optogenetic stimulation-activated neurons (expressing the Fos protein) and footshock-activated neurons (expressing the *fos* mRNA) (*Figure 6−figure supplement 1B−E*). Then we injected the AAV2/9-EF1a-DIO-ChR2-mCherry virus into the PBN and implanted the optoelectrode into the PVT of *VgluT2-ires-Cre* mice (*Figure 6A*). We first recorded the spiking signals in response to 10 sweeps of 2 s laser pulse trains (20 Hz, 5 mW, 5 ms). Then we recorded the spiking signals in response to 20 sweeps of 2 s footshock (0.5 mA) without laser in odd number sweeps or with laser in even number sweeps (*Figure 6A*). We found that laser or footshock (without laser) increased firing rates in 22 or 28 of 40 units (*Figure 6B−C*). And there was also a broad overlap between laser-activated and footshock-activated units (*Figure 6D*). It was consistent with the dual Fos staining result, suggesting that PVT_PBN_ neurons are activated by aversive stimulation. Next, we analyzed the firing rates of PVT neurons in footshock with laser sweeps and footshock without laser sweeps (*Figure 6E-G*). We found that the footshock stimulus with laser activated 30 of 40 units (*Figure 6H*) and increased the overall firing rates of neurons compared with the footshock without laser result (*Figure 6I*). These results indicate that activation of the PBN-PVT projection could enhance the PVT neuronal responses to aversive stimulation.

**Figure 6.**
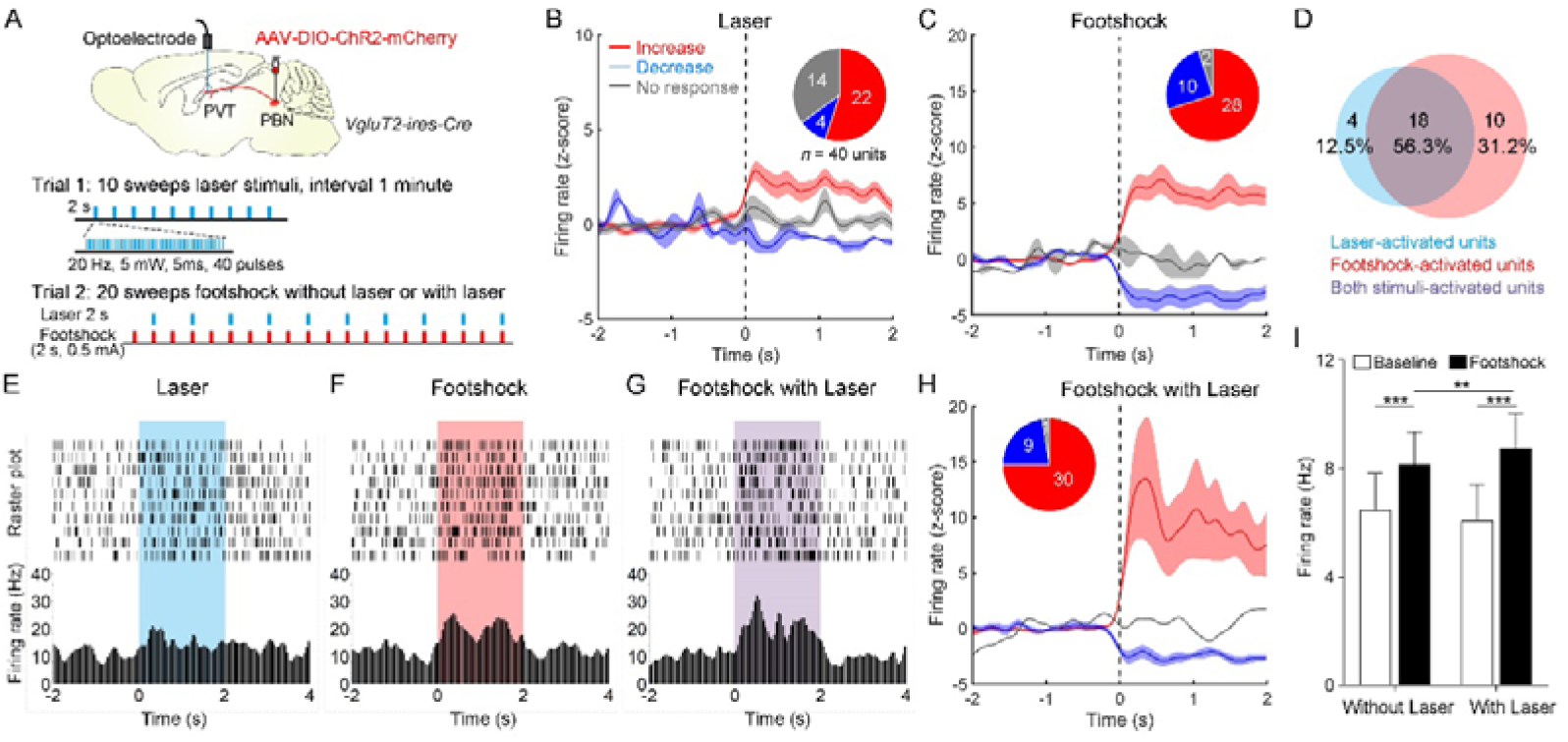
Neuronal activity of the PVT neurons in response to the footshock was modulated by the PBN-PVT projection. (A) Top: Schematic showed injection of AAV2/9-EF1a-DIO-ChR2-mCherry into the PBN and placement of the optoelectrode above the PVT of *VgluT2-ires-Cre* mice. Bottom: The protocol of 10 sweeps of laser stimuli (Trial 1) and 20 sweeps of footshock stimuli without or with laser (Trial 2). (B) Firing rates (z-score) of 40 units during laser stimuli (20 Hz, 5 mW, 5 ms, 2 s). Inserted: percentages of different groups of neurons according to z-score. (C) Firing rates (z-score) of 40 units during footshock (0.5 mA, 2 s) without laser stimuli. (D) Percentage of laser-activated, footshock-activated, and both stimuli-activated units. (E-G) Rastergrams and firing rates showed the spiking activity of one PVT neuron during laser stimulus (E), footshock without laser stimulus (F), and footshock with laser stimulus (G). (H) Firing rates (z-score) of 40 units during footshock (0.5 mA, 2 s) with laser stimuli (20 Hz, 5 mW, 5 ms, 2 s). (I) Quantification of the firing rates of 40 units before and during footshock without and with laser, *n* = 40 units. **p < 0.01, ***p < 0.001, all data were represented as mean ± SEM, two-way ANOVA followed by Bonferroni test for I. The following figure supplements are available for figure 6: **Figure 6−figure supplement 1.** Dual Fos staining detecting Fos protein and *fos* mRNA induced by laser stimulation and footshock.

### Pharmacogenetic activation of the PVT_PBN_ neurons induces anxiety-like behaviors

To further investigate the specific role of PVT_PBN_ neurons in modulating negative affective states, we injected the AAV2/1-hSyn-Cre virus into the bilateral PBN and injected the AAV2/9-hSyn-DIO-hM3Dq-mCherry virus or control virus into the PVT to activate the PVT_PBN_ neurons *(Figure 7A*). The majority of the PVT_PBN_ neurons could be activated by CNO in the hM3Dq expressing mice but not the control mice (*Figure 7B−D*). We found that pharmacogenetic activation of the PVT_PBN_ neurons reduced the center time (*Figure 7E−F*). Similarly, the time spent in open quadrants was decreased in the EZM of hM3Dq expressing mice (*Figure 7G−H*). And the unmoving time in the EZM test had an increased tendency in hM3Dq expressing mice (*Figure 7I*). We did not observe obvious nociception-related behaviors, such as forelimb wiping, hindlimb flinching, licking, or bitting during the experiments. These results indicate that pharmacogenetic activation of PVT_PBN_ neurons induces anxiety-like behaviors.

**Figure 7.**
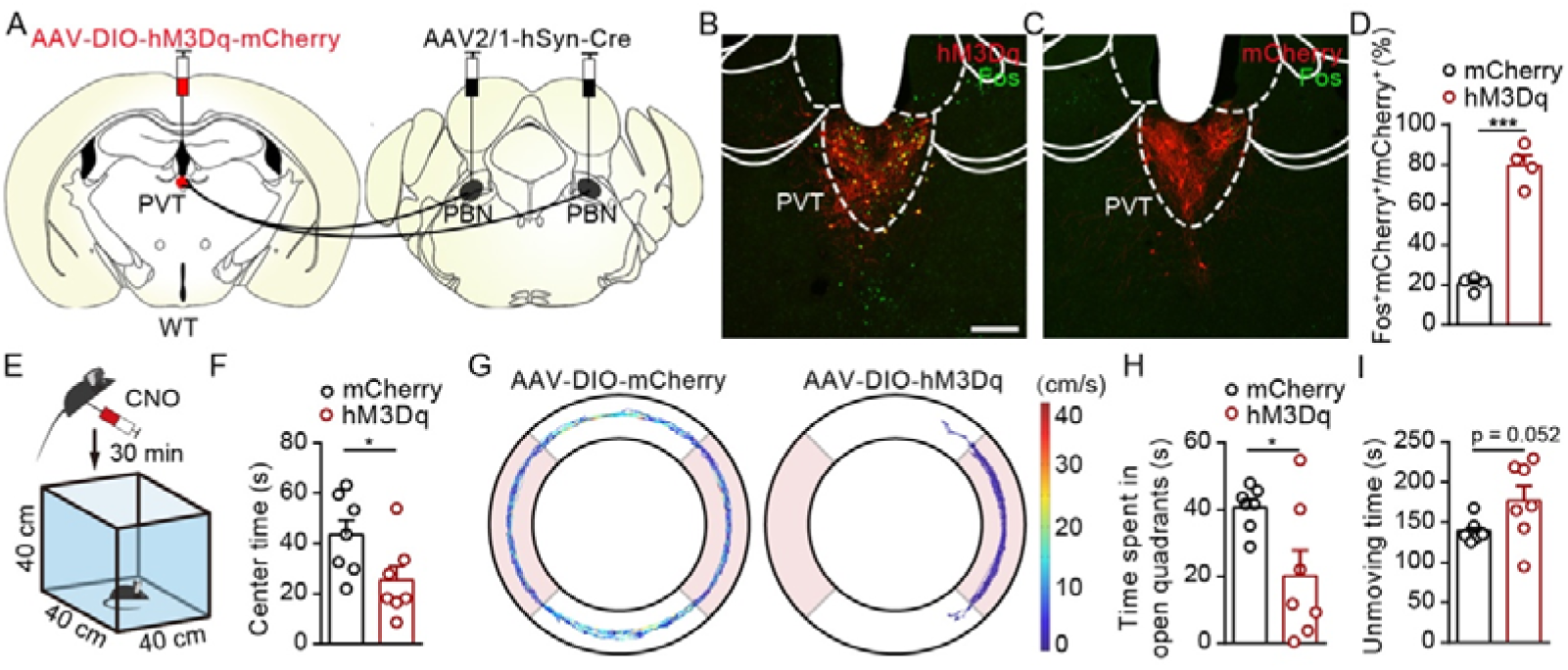
Activation of PVT_PBN_ neurons induced anxiety-like behaviors. (A) The illustration showed injection of AAV2/1-hSyn-Cre into the PBN and AAV2/9-hSyn-DIO-hM3Dq-mCherry into the PVT. (B and C) CNO administration evoked Fos expression in AAV2/9-hSyn-DIO-hM3Dq-mCherry injected mice (B) but not in AAV2/9-EF1a-DIO-mCherry injected mice (C). Scale bar: 200 μm. (D) Percentage of co-labeled neurons in the PVT, *n* = 4 mice per group. (E) The illustration of the OFT test with pharmacogenetic activation. (F) Quantification of center time in the OFT, *n* = 7 mice per group. (G) Example of elevated zero maze (EZM) traces from the mice injected with AAV2/9-EF1a-DIO-mCherry or AAV2/9-hSyn-DIO-hM3Dq-mCherry. (H and I) Quantification of the time spent in open quadrants (H) and the unmoving time in the EZM test (I), *n* = 7 mice per group. *p < 0.05, ***p < 0.001, all data were presented as mean ± SEM. Unpaired student’s *t*-test for D, F, H, and I. The following figure supplements are available for figure 7: **Figure 7−figure supplement 1.** Distribution pattern of projection fibers of PVT_PBN_ neurons.

Furthermore, we examined the anatomic distribution of terminals of PVT_PBN_ neurons. We labeled the PVT_PBN_ neurons in WT mice by injecting the AAV2/1-hSyn-Cre virus into the PBN and AAV2/8-EF1a-DIO-EGFP virus into the PVT (*Figure 7−figure supplement 1A−B*). We found that the PVT_PBN_ neurons sent projections to several brain areas, in particular the nucleus accumbens core (NAc), BNST, and CeA (*Figure 7−figure supplement 1B−H*), which was similar to the early tracing research of PVT efferent projections *(Kirouac, 2015)*.

## Discussion

In this study, we employed viral tracing and electrophysiology to confirm the monosynaptic excitatory connectivity between the PBN and the PVT. Optogenetic or pharmacogenetic activation of the PBN-PVT projection or the PVT-projecting PBN neurons induced anxiety-like, aversion-like, and fear-like behaviors. Optogenetic inhibition of the PBN-PVT projection or the PVT-projecting PBN neurons could partially reduce 2-MT induced aversive behaviors as well as footshock-induced freezing behaviors. The activity of PVT_PBN_ neurons was increased in several aversive stimuli and could be further increased by activation of PBN-PVT projection. Besides, activation of PVT_PBN_ neurons induced anxiety-like behaviors. Taken together, our results reveal the functional role of the PBN-PVT projection in modulating negative affective states in mice.

### PBN efferents and PBN-PVT monosynaptic excitatory projection

The PBN is a critical hub receiving sensory information from the spinal cord *(Todd, 2010)*. The widespread distribution of PBN efferents contributes to different aspects of behavioral and physiological responses. Previous studies showed that the CGRP-expressing neurons in the PBN project to the CeA contribute to the affective dimension of pain. In contrast, non-CGRP neurons may transmit sensory pain information *(Han et al., 2015)*. The projections from the PBN to the VMH or PAG are involved in producing escape behaviors to avoid injury, while the projections from the PBN to the BNST or CeA participate in facilitating aversive memory *(Chiang et al., 2020)*. The PBN neurons, which receive projections from the spinal cord, form strong functional synaptic connections with the ILN neurons but not the CeA neurons to process the nociceptive signals *(Deng et al., 2020)*. The PVT nucleus located in the middle line of the brain is an important area that participates in affective states processing *(Kirouac, 2015)*. Although recent research has reported that the projecting fibers from PBN were found in the PVT *(Chiang et al., 2020)*, remarkably little is known about the connectivity information and function of the PBN-PVT projection.

Since we injected the constitutively expressed ChR2 virus into the PBN, few neurons in the LC (which is medial to the PBN) might be infected. The LC neurons express the VgluT1 and also project to the PVT (*Beas et al., 2018*). Although the PBN-PVT projection comprises the major portion of the projections, there is still potential contamination from the LC-PVT projections. We also observed a small portion of inhibitory connections between the PBN and the PVT. It is consistent with the previous study showing GABAergic neurons in the PBN also send sparse projections to the PVT *(Chiang et al., 2020)*. Further in situ hybridization results confirmed that the PVT-projecting PBN neurons are mainly glutamatergic neurons expressing *Vglut2* mRNA. These results suggest that the majority of the PBN-PVT projection appeared to be excitatory.

We also found that the density of PBN glutamatergic is higher in the middle and posterior PVT (pPVT). These results are consistent with the various studies supporting that pPVT is a particularly aversive region of the PVT (*Gao et al., 2020*; *Beas et al., 2018*; *Barson et al., 2020*).

### PBN-PVT projection modulates negative affective states

We found that activation of PBN-PVT projection or the PVT-projecting PBN neurons induced anxiety-like behaviors and fear-like behaviors in the OFT and EZM. We observed that mice displayed robust running and jumping behaviors mainly in the first minute in optogenetic manipulation, and these phenomena were not observed in the pharmacogenetic experiment. These might be caused by an instantly increased activity of PBN-PVT projection induced by optogenetic manipulation. Mice might display “fight or flight” during sudden affective state transitions. And the pharmacogenetic approach takes several minutes to gradually enhance neural activity, and the affective state changes in a relatively mild way. We also observed that the anxiety-like behaviors in the OFT still existed several minutes after optogenetic activation of the PBN-PVT projection. However, in the RTPA test, the aversion appeared when the laser was on and disappeared when the laser was off, indicating the aversion was transient and could not be translated to associative learning. It was further confirmed by the prolonged condition place aversion test and cue-dependent optogenetic conditioning test. These results suggest that activation of the PBN-PVT induces instant negative affective states but does not drive associative fear memory formation.

Selectively optogenetic inhibition of PBN-PVT projection or PVT-projecting PBN neurons could reduce aversion-like and fear-like behaviors. To better examine the behavioral changes, we performed 10 minutes test in 2-MT and footshock experiments. So we used a relatively long-term protocol in optogenetic inhibition experiments (10 minutes constant laser). Such long-term inhibition protocols were used in other studies (*Zhou et al., 2019*; *Sun et al., 2020*). We also performed the classical fear conditioning test and found that inhibition of the PBN-PVT projection did not affect associative fear memory formation or retrieval, suggesting that the PBN-PVT projection mainly promotes aversion but does not facilitate negative association.

Our calcium imaging and Fos staining results indicated PVT neurons were activated after exposure to aversive stimuli, consistent with a previous study *(Zhu et al., 2018)*. The dual Fos staining experiment and optoelectrode experiments confirmed a broad overlap between laser-activated and footshock-activated neurons. Further analysis showed that activation of the PBN-PVT projection enhanced the overall firing rates of PVT neurons in response to footshock. These results suggest that the activation of the PBN-PVT could enhance the neuronal activity in response to aversive stimulation.

The activation of PBN innervated PVT neurons induced anxiety-like behaviors, suggesting the PVT_PBN_ neurons are involved in modulating negative affective states. Previous studies have reported that activation of the PBN-CeA pathway is sufficient to drive a series of negative affective states behaviors *(Bowen et al., 2020; Cai et al., 2018; Han et al., 2015)*, enable associative learning, and generate aversive memory *(Chiang et al., 2020)*. Distinct from the PBN-CeA projection, we found that activation of the PBN-PVT projection only induced transient aversion-related behaviors, and inhibition of the PBN-PVT projection did not affect fear memory acquisition or retrieval. A study reported that only a few Fluoro-gold (FG)/tetramethylrhodamine-dextran (TMR) double-labeled neurons were sparsely distributed in the PBN of the mice injected with FG into the PVT and TMR into the CeA *(Liang et al., 2016)*. Our results also showed few collateral projecting fibers in the CeA or VMH from the PVT-projecting PBN neurons. These results suggested that the PBN-PVT pathway and the PBN-CeA pathway are two parallel pathways originating from distinct efferent neurons within the PBN to perform distinct functions. However, we also observed collateral projection fibers in BNST, LH, PVN, PAG, but not CEA or VMH. The PBN-PAG projection is suggested to mediate escaping behaviors *(Chiang et al., 2020)*. The possibility of antidromic effects following photoactivation of PBN terminals in PVT should be reminded.

The tracing results showed the PVT_PBN_ neurons projected to multiple brain areas, particularly the NAc, BNST, and CeA. The BNST and CeA have been previously implicated in negative affective behaviors *(Jennings et al., 2013; Tye et al., 2011)*. Previous studies showed that the activation of the PVT-CeA projections induces place aversion, and the effect persists on the next day in the absence of photostimulation (*Do Monte et al., 2017*). Similarly, long-term depression (LTD)-like the stimulation of PVT-CeA projections or inhibition of the same circuit induces a persistent attenuation of fear responses (*Chen and Bi., 2019*; *Do Monte et al., 2015*; *Penzo et al., 2015*). These results revealed a critical role of the PVT-CeA projection in aversive memory formation. In our study, we found that PBN-PVT is not crucial for aversive memory formation. The possible reason might be that manipulation of the PVT-CeA induces direct excitatory inputs to the CeA, and the inputs are strong enough for aversive memory formation. However, activation of the PBN-PVT projection might not induce enough excitatory inputs to the CeA via the disynaptic connection.

A study also found that PVT mediates descending pain facilitation underlying persistent pain conditions via the PVT-CeA-PAG circuit *(Liang et al., 2020)*. Different downstream pathways of PVT_PBN_ neurons might have different functions and deciphering the circuit mechanisms needs further examination.

### The potential role of PBN-PVT projection in depression and pain

It is worth noting that although the pharmacogenetic activation of the PVT-projecting PBN neurons induced anxiety-like behaviors and fear-like behaviors in the hM3Dq group mice, no depression-like symptoms were observed in TST and FST. On the other side, chronic pain models, such as the partial sciatic nerve ligation model, the spared nerve injury model, and complete Freund’s adjuvant model, generally induce anxiety and depression at least 3−4 weeks after the surgery in mice *(Dimitrov et al., 2014; Zhou et al., 2019)*. Our study collected the behavioral data 30 minutes after a single dose of CNO injection. Different behavioral tests were performed at least three days apart to eliminate the residual CNO effects. We hypothesized that depression-like behaviors might be observed if we repeatedly activate the PBN-PVT projection for weeks. However, whether the PBN-PVT is involved in depression is still unknown.

A recent study revealed that the PBN neurons convey nociception information from the spinal cord to the ILN, which is relatively closed to the PVT *(Deng et al., 2020)*. In our results, we carefully checked the virus expression and optic fiber locations of mice. We found that pharmacogenetic activation of PVT-projecting PBN neurons did not affect the basal nociceptive thresholds or formalin-induced licking behaviors. Moreover, no obvious nociception-related behaviors (such as forelimb wiping, hindlimb flinching, licking, or biting) were found through specific manipulation of the PBN innervated PVT neurons, which suggests that the PBN-PVT projection might be not involved in nociceptive information processing.

In summary, we identified the functional role of the PBN-PVT projection in modulating negative affective states. Our study paves the way for further deciphering the distinct roles of the PBN neural circuit in affective behaviors.

## Acknowledgements

We thank Dr. Hua-Tai Xu for providing *Rosa26-tdTomato* mice. We thank Dr. Yan-Gang Sun for providing *VgluT2-ires-Cre* mice. We thank all the lab members of D.M. for their helpful discussion. This work was supported by the National Natural Science Foundation of China (No. 31900717, 31571086), the Shanghai Sailing Program (19YF1438700 to D.M.), the Young Elite Scientists Sponsorship Program of China Association for Science and Technology (2019QNRC001 To D.M.), and the Starting Research Fund from the Shanghai General Hospital.

## Author contributions

Y.B.Z. performed the virus injection experiments and behavioral experiments. Y.W. performed the dual Fos staining experiments. X.X.H. and Y.W. performed the optoelectrode experiments. R.Z. performed the electrophysiological experiments. Y.B.Z., Y.W., M.Z.L., and P.F.L. performed the histological experiments. J.B.L., L.Z., and D.M. designed the experiments. Y.B.Z. and D.M. wrote the manuscript.

## Materials and methods

### Animals

Male C57Bl/6J wild-type mice, *Rosa26-tdTomato* mice (Jax Stock No: 007909, gifted from Dr. Hua-Tai Xu, Institutes of Neuroscience, Chinese Academic of Sciences), *VgluT2-ires-Cre* mice (Jax Stock No: 016963, gifted from Dr. Yan-Gang Sun, Institutes of Neuroscience, Chinese Academic of Sciences) were used. Animals were housed in standard laboratory cages in a temperature (23-25°C)-controlled vivarium with a 12:12 light/dark cycle, free to food and water. For tracing and behavioral experiments, the mice were injected with the virus at 7−8 weeks old and performed the behavioral tests at 11−12 weeks old. For the electrophysiological experiments, the mice were injected with the virus at 4−6 weeks old to accomplish the electrophysiological experiments at 7−9 weeks old. For *in vivo* fiber photometry and optoelectrode experiments, the mice were injected with the virus at 7−8 weeks old to accomplish the experiments at 10−11 weeks old. All animal experiment procedures were approved by the Animal Care and Use Committee of Shanghai General Hospital (2019AW008).

### Stereotaxic surgery

Mice were anesthetized by vaporized sevoflurane (induction, 3%; maintenance, 1.5%) and head-fixed in a mouse stereotaxic apparatus (RWD Life Science Co.).

For electrophysiological experiments, the AAV2/8-hSyn-ChR2-mCherry virus (300 nl, 4 x 10^12^ v.g./ml, AG26976, Obio Technology) was injected into the PBN nucleus of WT mice in the stereotaxic coordinate: anteroposterior (AP) −5.2 mm, mediolateral (ML) +1.3 mm, and dorsoventral (DV) −3.4 mm.

For tracing studies, the AAV2/8-EF1a-DIO-EGFP virus (300 nl, S0270, Taitool Bioscience) was injected into the PBN (mentioned above) of *VgluT2-ires-Cre* mice.

For the retrovirus injection surgery, the retrograde transport Cre recombinase retroAAV2/2-hSyn-Cre virus (150 nl, 4 x 10^12^ v.g./ml, S0278-2RP-H20, Taitool Bioscience) was injected in the *Rosa26-tdTomato* mice at two locations of PVT respectively: (1) AP −1.22 mm, ML 0 mm, DV −2.9 mm; (2) AP −1.46 mm, ML 0 mm, DV −2.9 mm.

For optogenetic activation of PVT-projecting PBN fibers, the AAV2/9-EF1a-DIO-ChR2-mCherry virus (300 nl, 4 x 10^12^ v.g./ml, S0170, Taitool Bioscience) or the AAV2/9-EF1a-DIO-mCherry virus (300 nl, 4 x 10^12^ v.g./ml, AG20299, Obio Technology) were bilaterally injected into the PBN (mentioned above) of *VgluT2-ires-Cre* mice, and a 200 μm diameter optic fiber was implanted over the PVT (AP −1.46 mm, ML 0 mm, DV −2.9 mm) with a 20° angle towards the midline.

For the pharmacogenetic activation of PVT-projecting PBN neurons, the retroAAV2/2-hSyn-Cre virus (150 nl, 4 x 10^12^ v.g./ml, S0278-2RP-H20, Taitool Bioscience) was injected into the PVT (AP −1.46 mm, ML 0 mm, DV −2.9 mm), the AAV2/9-hSyn-DIO-hM3Dq-mCherry virus (300 nl, 4 x 10^12^ v.g./ml, PT-0019, BrainVTA) or the control AAV2/9-EF1a-DIO-mCherry virus were bilateral injected into the PBN (mentioned above) of the WT mice.

For optogenetic inhibition of PVT-projecting PBN fibers, AAV2/9-EF1a-DIO-NpHR3.0-EYFP virus (300 nl, 4 x 10^12^ v.g./ml, AG26966, Obio Technology) or the AAV2/8-EF1a-DIO-EGFP virus were bilaterally injected into the PBN (mentioned above) of *VgluT2-ires-Cre* mice, and a 200 μm diameter optic fiber was implanted over the PVT (AP −1.46 mm, ML 0 mm, DV −2.9 mm) with a 20° angle towards the midline.

For optogenetic inhibition of PVT-projecting PBN neurons, retroAAV2/2-hSyn-Cre was injected into the PVT(AP −1.46 mm, ML 0 mm, DV −2.9 mm), AAV2/9-EF1a-DIO-NpHR3.0-EYFP virus (300 nl, 4 x 10^12^ v.g./ml, AG26966, Obio Technology) or the AAV2/8-EF1a-DIO-EGFP virus was injected into the PBN of WT mice, the left optic fiber was implanted over the PBN vertically and the right one were placed over the PBN with a 20° angle towards the midline.

For *in vivo* fiber photometry experiments, the AAV2/8-hSyn-GCaMP6s virus (200 nl, 4 x 10^12^ v.g./ml, S0225-8, Taitool Bioscience) was injected into the PVT nucleus (AP −1.46 mm, ML 0 mm, DV −2.90 mm) of the WT mice, the optic fiber was implanted above the PVT with a 20° angle towards the midline.

For optoelectrode experiments, the AAV2/9-EF1a-DIO-ChR2-mCherry virus (300 nl, 4 x 10^12^ v.g./ml, AAV2/9-S0170, Taitool Bioscience) were bilaterally injected into the PBN (mentioned above) of *VgluT2-ires-Cre* mice. Three weeks later, the homemade optoelectrode was implanted into the PVT nucleus (AP −1.46 mm, ML 0 mm, DV −2.90 mm).

For pharmacogenetic activation of PVT_PBN_ neurons, the AAV2/1-hSyn-Cre virus (300 nl, 1.5 x 10^13^ v.g./ml, S0278-1-H50, Taitool Bioscience) was bilaterally injected into the PBN nucleus, the AAV2/9-hSyn-DIO-hM3Dq-mCherry virus or the control AAV2/9-EF1a-DIO-mCherry virus was injected into the PVT (AP −1.46 mm, ML 0 mm, DV −2.9 mm) of the WT mice.

The virus was infused through a glass pipette (10−20 μm in diameter at the tip) at the rate of 50−100 nl/minute. The injection pipette was left in place for additional 8 minutes. After the surgeries, the skin was closed by the sutures, and the optic fiber was secured through the dental acrylic. Generally, tracing, electrophysiological or behavioral experiments were performed at least three weeks later. After experiments, histological analysis was used to verify the location of viral transduction and the optic fiber. The mice without correct transduction of virus or correct site of optic fiber were excluded for analysis.

### Histology

Animals were deeply anesthetized with vaporized sevoflurane and transcardially perfused with 20 ml saline, followed by 20 ml paraformaldehyde (PFA, 4% in PBS). Brains were extracted and soaked in 4% PFA at 4°C for a minimum of 4 hours and subsequently cryoprotected by transferring to a 30% sucrose solution (4°C, dissolved in PBS) until brains were saturated (for 36−48 hours). Coronal brain sections (40 μm) were cut using a freezing microtome (CM1950, Leica). The slices were collected and stored in PBS at 4°C until immunohistochemical processing. Nuclei were stained with DAPI (Beyotime, 1:10000) and washed three times with PBS.

The brain sections undergoing immunohistochemical staining were washed in PBS 3 times (10 minutes each time) and incubated in a blocking solution containing 0.3% TritonX-100 and 5% normal donkey serum (Jackson ImmunoResearch, USA) in PBS for 1 hour at 37°C. Sections were then incubated (4°C, 24 hours) with primary antibodies dissolved in 1% normal donkey serum solution. Afterward, sections were washed in PBS 4 times (15 minutes each time), then incubated with secondary antibodies for 2 hours at room temperature. After DAPI staining and washing with PBS, sections were mounted on glass microscope slides, dried, and covered with 50% glycerin (ThermoFisher). The images were taken by the Leica DMi8 microscope and the Leica SP8 confocal microscopy. The images were further processed by Fiji and Photoshop.

### RNAscope in situ hybridization

Mice were anesthetized with isoflurane and rapidly decapitated. Brains were roughly dissected from perfused mice and post-fixed in 4% PFA at 4 **°**C overnight, dehydrated in 30% sucrose 1×PBS at 4 °C for 2 days. Mouse brains were embedded in OCT compound, cryosectioned in 15 µm coronal slices, and mounted on SuperFrost Plus Gold slides (Fisher Scientific). In situ hybridization was performed according to the protocol of the RNAscope Multiplex Fluorescent Reagent Kit v2 (Cat. No. 320293). Probes were purchased from Advanced Cell Diagnostics: *c-Fos* (Cat. No. 316921-C2), *Tac1* (Cat. No. 410351-C2), *Tacr1* (Cat. No. 428781-C2), *Pdyn* (Cat. No. 318771), and *VgluT2* (Cat. No. 319171-C2). Primary antibodies include rabbit anti-c-Fos (Abcam, cat. No. ab190289, 1:4000), goat anti-CGRP (Abcam, ab36001, 1:1000), and rabbit anti-DsRed (Clontech, Cat. No. 632496, 1:500). All secondary antibodies were purchased from Jackson ImmunoResearch and used at 1:400 dilution. Secondary antibodies include Alexa 488 donkey anti-rabbit (Cat. No. 711-545-152), Cy3 donkey anti-rabbit (Cat. No. 711-165-152), and Alexa 488 donkey anti-goat (Cat. No. 705-546-147). Images were collected on a Leica fluorescence microscope and Leica LAS Software.

### Fos induction

The mice were habituated for three days and performed gentle grabbing and holding for 1 minute, five times every day to minimize background Fos expression.

To study the effect of pharmacogenetic manipulations on PVT-projecting PBN neurons, we intraperitoneally injected 0.5 mg/kg clozapine N-oxide (CNO, Sigma). Ninety minutes later, the brain tissues were processed. To assess 2-MT evoked Fos expression in the PVT, the mice were kept in a chamber with a floor covered with cotton containing 100 ml, 1:1000 2-MT volatilized the predator odor for 90 minutes. To assess footshock-induced Fos expression in the PVT, we placed the mice into the chamber and delivered 30 times inevitable footshock (0.5 mA, 1 second) with a variable interval (averaging 60 seconds). After stimulation, animals were kept in the same apparatus for another 60 minutes, and brain tissues were then processed.

For the dual Fos experiments, we first delivered 20 minutes 473 nm laser pulses (20 Hz, 5 mW, 5ms) and left the mice to rest in the homecage for 60 minutes. Then we delivered the 20 minutes footshock stimulus (0.5 mA, 1 s, 30 times) and perfused the mice.

### Electrophysiology

The electrophysiological experiment was performed as previously described *(Mu et al., 2017)*. Mice were anesthetized with sevoflurane and perfused by the ice-cold solution containing (in mM) sucrose 213, KCl 2.5, NaH_2_PO_4_ 1.25, MgSO_4_ 10, CaCl_2_ 0.5, NaHCO_3_ 26, glucose 11 (300−305 mOsm). Brains were quickly dissected, and the coronal slice (250 μm) containing the PBN or PVT were chilled in ice-cold dissection buffer using a vibratome (V1200S, Leica) at a speed of 0.12 mm/second. The coronal sections were subsequently transferred to a chamber and incubated in the artificial cerebrospinal fluid (ACSF, 34°C) containing (in mM): NaCl 126, KCl 2.5, NaH_2_PO_4_ 1.25, MgCl_2_ 2, CaCl_2_ 2, NaHCO_3_ 26, glucose 10 (300−305 mOsm) to recover for at least 40 minutes, then kept at room temperature before recording. All solutions were continuously bubbled with 95% O_2_/5% CO_2_.

All experiments were performed at near-physiological temperatures (30−32°C) using an in-line heater (Warner Instruments) while perfusing the recording chamber with ACSF at 3 ml/minute using a pump (HL-1, Shanghai Huxi). Whole-cell patch-clamp recordings were made from the target neurons under IR-DIC visualization and a CCD camera (Retiga ELECTRO, QIMAGING) using a fluorescent Olympus BX51WI microscope. Recording pipettes (2−5 MΩ; Borosilicate Glass BF 150-86-10; Sutter Instrument) were prepared by a micropipette puller (P97; Sutter Instrument) and backfilled with potassium-based internal solution containing (in mM) K-gluconate 130, MgCl_2_ 1, CaCl_2_ 1, KCl 1, HEPES 10, EGTA 11, Mg-ATP 2, Na-GTP 0.3 (pH 7.3, 290 mOsm) or cesium-based internal solution contained (in mM) CsMeSO_3_ 130, MgCl_2_ 1, CaCl_2_ 1, HEPES 10, QX-314 2, EGTA 11, Mg-ATP 2, Na-GTP 0.3 (pH 7.3, 295 mOsm). Biocytin (0.2%) was included in the internal solution.

In PBN-PVT ChR2 experiments, whole-cell recordings of PBN neurons with current-clamp (I = 0 pA) were obtained with pipettes filled with the potassium-based internal solution. The 473 nm laser (5 Hz, 10 Hz, 20 Hz pulses, 0.5 ms duration, 2 mW/mm^2^) was used to activate PBN ChR2 positive neurons. Light-evoked EPSCs and IPSCs of PVT neurons recorded with voltage-clamp (holding voltage of -70 mV or 0 mV) were obtained with pipettes filled with the cesium-based internal solution. The 473 nm laser (20 Hz paired pulses, 1 ms duration, 4 mW/mm^2^) was used to activate ChR2 positive fibers. The light-evoked EPSCs were completely blocked by 1 μM TTX (tetrodotoxin), rescued by 100 μM 4-AP (4-Aminopyridine), and blocked by 10 μM NBQX (6-nitro-7-sulphamoylbenzo(f)quinoxaline-2,3-dione). NBQX and TTX were purchased from Tocris Bioscience. All other chemicals were obtained from Sigma.

Voltage-clamp and current-clamp recordings were carried out using a computer-controlled amplifier (MultiClamp 700B; Molecular Devices, USA). During recordings, traces were low-pass filtered at 4 kHz and digitized at 10 kHz (DigiData 1550B1; Molecular Devices). Data were acquired by Clampex 10.6 and filtered using a low-pass-Gaussian algorithm (-3 dB cut-off frequency = 1000 Hz) in Clampfit 10.6 (Molecular Devices).

### Optogenetic manipulation

For activating the PBN-PVT projection, a 473 nm laser (20 Hz, 5 ms pulse duration, 5 mW) was delivered. For inhibition of the PBN-PVT projection and the PVT-projecting PBN neurons, a constant laser (589 nm, 10 mW) was delivered.

### Pharmacogenetic manipulation

All behavioral tests were performed 30 minutes after intraperitoneal injection of 0.5 mg/kg CNO in pharmacogenetic manipulation. Different behavior tests were performed at least three days apart.

### Open field test

The open field test (OFT) was used to assess locomotor activity and anxiety-related behavior in an open field arena (40 x 40 x 60 cm) with opaque plexiglass walls. The mouse was placed in the center of the box and recorded by a camera attached to a computer. The movement was automatically tracked and analyzed by AniLab software (Ningbo AnLai, China). The total distance traveled, the total velocity, the total unmoving time (the mice were considered to be unmoving if unmoving time lasts more than 1 s), and time spent in the center area (20 x 20 cm) were measured. The box was cleaned with 70% ethanol after each trial.

To assess the effect of optogenetic activation of the PBN-PVT projection, 15 minutes sessions consisting of 5 minutes pre-test (laser OFF), 5 minutes laser on test (laser ON), and 5 minutes post-test (laser OFF) periods. Laser (473 nm, 20 Hz, 5 ms, 5 mW) was delivered during the laser on phase.

To assess the effect of pharmacogenetic manipulations of PVT-projecting PBN neurons on locomotor activity and affective behaviors, we recorded the the movement 30 minutes after intraperitoneal (i.p.) injection with CNO.

To assess the effect of inhibition of the PBN-PVT projection on the aversive behaviors induced by 2-MT. One cotton ball containing 5 ml 2-MT (1:1000) solution was placed on the center of the upper left quadrant to disseminate fear-odor, then a constant laser (589 nm, 10 mW) was delivered during the 10 minutes test. The time spent in the 2-MT paired quadrant was calculated.

### Elevated zero maze (EZM)

The EZM was an opaque plastic circle (60 cm diameter), which consisted of four sections with two opened and two closed quadrants. Each quadrant had a path width of 6 cm. The maze was elevated 50 cm above the floor. The animals were placed into an open section facing a closed quadrant and freely explored the maze for 5 minutes.

### Real-time place aversion (RTPA) test

Mice were habituated to a custom-made 20 x 30 x 40 cm two-chamber apparatus (distinct wall colors and stripe patterns) before the test. Each mouse was placed in the center and allowed to explore both chambers without laser stimulation for 10 minutes on Day 1. The movement was recorded for 10 minutes as a baseline. The mice performed a slight preference for the black chamber according to the fact the mice have innate aversive to brightly illuminated areas. On Day 2, 473 nm laser stimulation (20 Hz, 5 ms, 5 mW) was automatically delivered when the mouse entered or stayed in the black chamber and turned off when the mouse exited the black chamber for 10 minutes. Finally, the mouse was allowed to freely explore both chambers without laser stimulation for another 10 minutes. The RTPA location plots and total time on the stimulated side were recorded and counted with the AniLab software.

### Condition place aversion (CPA)

After habituation, mice were placed in the center of the two-chamber apparatus and allowed to explore either chamber for 15 minutes on Day 1. On Day 2, mice were restricted to one chamber (laser paired chamber) with photostimulation (473 nm, 20 Hz, 5 ms, 5 mW) for 30 minutes in the morning and restricted to the other chamber (unpaired chamber) without photostimulation in the afternoon. On Day 3, mice were restricted to the unpaired chamber without photostimulation in the morning and restricted to the laser paired chamebr with photostimulation in the afternoon. On Day 4, mice were allowed to explore both chambers without laser stimulation for another 15 minutes. The time in the laser-paired chamber was calculated on Day 1 and Day 4.

### 2-MT-induced aversion

To assess the effect of optogenetic inhibition of the PBN-PVT projection or PVT-projecting PBN neurons on the aversive state, three cotton balls containing 15 ml 2-MT (1:1000) solution were placed in the black chamber. A constant laser (589 nm, 10 mW) was delivered during the 10 minutes test.

### Cue-dependent optogenetic conditioning test

Video Freeze fear conditioning system with optogenetic equipment (MED Associates, MED-VFC-OPTO-USB-M) and Video Freeze software were used.

On Day 1, mice were habituated to the fear conditioning chambers and allowed to explore for 2 minutes freely, then three tones (75 dB, 4 kHz, 30 seconds duration) separated by a variable interval with a range of 60−120 seconds and the average of 90 seconds were delivered.

On Day 2, mice were trained with the sound cue (75 dB, 4 kHz, 30 seconds) paired with a simultaneous 30 seconds laser pulse train (20 Hz, 5 ms, 5 mW) for six times separated by a variable interval (averaging 90 seconds). The mice were kept in the conditioning chamber for another 60 seconds before returning to the home cages.

On Day 3, mice were placed back into the original training chamber for 3 minutes to perform the contextual test. After 2−3 hours, the conditioning chamber was modified by changing its metal floor and sidewalls. Mice were placed in the altered chamber for 3 minutes to measure the freezing level in the altered context. A tone (75 dB, 4 kHz) was delivered for 30 seconds to perform the cue test.

The behavior of the mice was recorded and analyzed with the Video Freeze software. Freezing was defined as the complete absence of movement for at least 0.5 seconds. On the conditioning day, the freezing percentages were calculated for 30 seconds after each tone/laser stimulus. For the contextual test, the freezing percentages were calculated for three minutes. For the cue test, the freezing percentages were calculated for 30 seconds during tone.

### Auditory fear conditioning test

On Day one, mice were habituated to the fear conditioning chambers. On Day two, mice were conditioned by seven trials of sound tone (75 dB, 4 kHz, 30 s) co-terminated with footshock (0.6 mA, 2 s) averagely separated by 90 seconds. Laser (589 nm, 10 mW) was delivered 1 second before the footshock and lasted for 4 seconds at each trial. On Day 3, mice were placed back into the original training chamber for 3 minutes to perform the contextual test, and the laser was delivered during the second minute. After 2−3 hours, the mice were placed into a modified chamber to perform the cue test. Three tones were given averagely separated by 90 seconds. The laser was delivered during the second tone.

The behavior of the mice was recorded and analyzed with the Video Freeze software. The freezing percentages of the 27 seconds tone before laser (to avoid the influence of laser) for each trial were summarized to indicate fear memory acquisition in the conditioning test. For the contextual test, the freezing percentages were calculated for every minute. For the cue test, the freezing percentages were calculated for 30 seconds during tone.

### Freezing behavior

For analyses of freezing behavior induced by pharmacogenetic activation of PVT-projecting PBN neurons, we injected CNO and recorded the mouse behavior using the Video Freeze fear conditioning system 30 minutes later.

The Video Freeze fear conditioning system (MED Associates, MED-VFC-OPTO-USB-M) was also used to assess the effect of optogenetic inhibition of PBN-PVT projection and the PVT-projecting PBN neurons on the fear-like behavior induced by footshock. After free exploration of the chamber for 2 minutes, 15 times footshocks (0.6 mA,1 second) were delivered within 10 minutes with a constant 589 nm laser (10 mW). The freezing percentages during 10 minutes were analyzed.

The Video Freeze fear conditioning system was also used to assess the effect of optogenetic inhibition of the PVT-projecting PBN neurons on the fear-like behavior induced by 2-MT. 10 ml 2-MT (1:1000) dissolved in the ddH_2_O was soaked into the cotton ball on the bottom of the training box. A constant laser (589 nm, 10 mW) was delivered during the tests.

### Tail suspension test (TST)

Mice were individually suspended by an adhesive tape placed roughly 2 cm from the tip of the tail and videotaped for 6 minutes. Mice were considered immobile without initiated movements, and the immobility time was scored in the last 3 minutes by an observer unknown of the treatments.

### Forced swim test (FST)

Mice were individually placed for 6 minutes in clear cylinders (45 cm height, 20 cm internal diameter) containing freshwater (25°C, 15 cm depth). The swimming activity was videotaped, and immobility time in the last 3 minutes was counted manually by an investigator unaware of animal grouping. The mice were considered immobile when they stopped swimming/struggling or only slightly moved to keep the nose above the surface.

### von Frey test

The von Frey test was used to assess the mechanical sensitivity *(Mu et al., 2017)*. The mice were acclimated to the observation chambers for two days (2 hours for each day) before the test. A series of von Frey hairs with logarithmically incrementing stiffness (0.16−2.0 grams) were used to stimulate the mouse hind paw perpendicularly. The 50% paw withdrawal threshold was determined using the up-down method.

### Hargreaves test

Hargreaves tests were performed as described previously *(Mu et al., 2017)*. Mice were placed in an individual plexiglass box with a glass floor. A radiant heat beam was exposed directly to the hind paw until the paw was withdrawn. The trials were repeated three times with an interval of at least 15 minutes. To avoid potential damage, the test was executed with a 20 seconds cut-off time.

### Formalin test

In the formalin test, the mice received an intraplantar injection of formalin (5%, 20 μl/mouse) and were placed into a plexiglass box (width: 10 cm, length: 10 cm, height: 15 cm) individually to record the pain-related licking behaviors for 1 hour. All videos were analyzed by trained investigators blinded to the experimental treatment of the animals.

### Rotarod test

Mice were trained twice on a rotarod apparatus (MED Associates) with a rod accelerated 5−20 revolutions per minute (r.p.m.) for 5 minutes before the experimental day. On the second day, each mouse underwent three trials with a rod was programmed to accelerate from 0 to 40 rpm over 300 seconds, then the average rpm at the point of falling was recorded.

### Fiber photometry

*In vivo* fiber photometry experiments were performed as previously described *(Zhu et al., 2020)*. After two weeks for virus expression, the mice were gently handled to be familiar with the calcium signal recording experiments (Thinker-Biotech). A signal (for synchronization) was manually tagged with the shock and air puff to evaluate the activity of PVT neurons. The calcium transient was recorded at 50 Hz. The fluorescence values change (ΔF/F) was calculated from the formula of (F−F0)/F0 and the F0 represented the median of the fluorescence values in the baseline period (−1 to −0.5 seconds relative to the stimulation onset). To precisely quantify the change of the fluorescence values across the shock or air-puff stimulation, we defined 0.5 to 1.0 seconds after the onset as the post-stimulus period.

### Optoelectrode recording and analysis

The homemade optoelectrode consisted of an optic fiber (200 mm in diameter) glued to 16 individually insulated nichrome wires (35 μm internal diameter, 300-900 Kohm impedance, Stablohm 675, California Fine Wire). The 16 microwire arrays were arranged in a 4 + 4 + 4 + 4 pattern and soldered to an 18-pin connector (Mil-Max). Three weeks after virus injection, the optoelectrode was implanted to the PVT nucleus (AP −1.46 mm, ML 0 mm, DV −2.90 mm). After one week of recovery, two trials were performed continuously. Trial 1 contained ten sweeps of 2 s laser pulse trains (473 nm, 5 ms, 20 Hz, 8 mW). The interval of sweeps was 60 s. Trial 2 contained twenty sweeps of 2 s footshock (0.5 mA). The interval of sweeps was 60 s. In the even time sweeps (2, 4, 6, 8,10, 12, 14, 16, 18, 20), 2 s laser pulse trains were delivered spontaneously with the 2 s footshock. Neuronal signals were recorded using a Zeus system (Zeus, Bio-Signal Technologies: McKinney, TX, USA), and Spike signals were filtered online at 300 Hz. At the end of the experiment, all animals were perfused to confirm the optical fiber sites. Only the data of animals with correct optical fiber sites and virus expression regions were analyzed.

The spikes were sorted by the valley-seeking method with Offline Sorter software (Plexon, USA) and analyzed with NeuroExplorer (Nex Technologies: Boston, MA, U.S.A.). Firing rates of the neurons and timestamps were exported for further analysis using customized scripts in MATLAB. The Kolmogorov-Smirnov (K-S) test was used to compare the spike firing rate of PVT during 2 s baseline (before stimulus) and 2 s after each stimulus. p < 0.001 indicated statistical significance. Z-score normalization maps were constructed from normalized firing rates.

### Quantification of the fiber intensity

For quantification of fluorescence of PVT_PBN_ efferents, the downstream targets of PVT_PBN_ neurons were taken photofluorograms with identical exposure time, the mean fluorescence value in each ROI (400 x 400 pixels) of each brain region was analyzed by Fiji. The fiber intensity was calculated as the fluorescence value of each brain region divided by that of the NAc. All data came from at least three different mice and were presented as mean ± SEM.

### Analysis

Statistical detection methods include unpaired student’s *t*-test, paired student’s *t*-test, one-way ANOVA with Bonferroni’s correction for multiple comparisons, two-way ANOVA with Bonferroni’s correction for multiple comparisons. A value of p < 0.05 was considered statistically significant. All data were represented as mean ± SEM.

---

**Figure 1−figure supplement 1.**
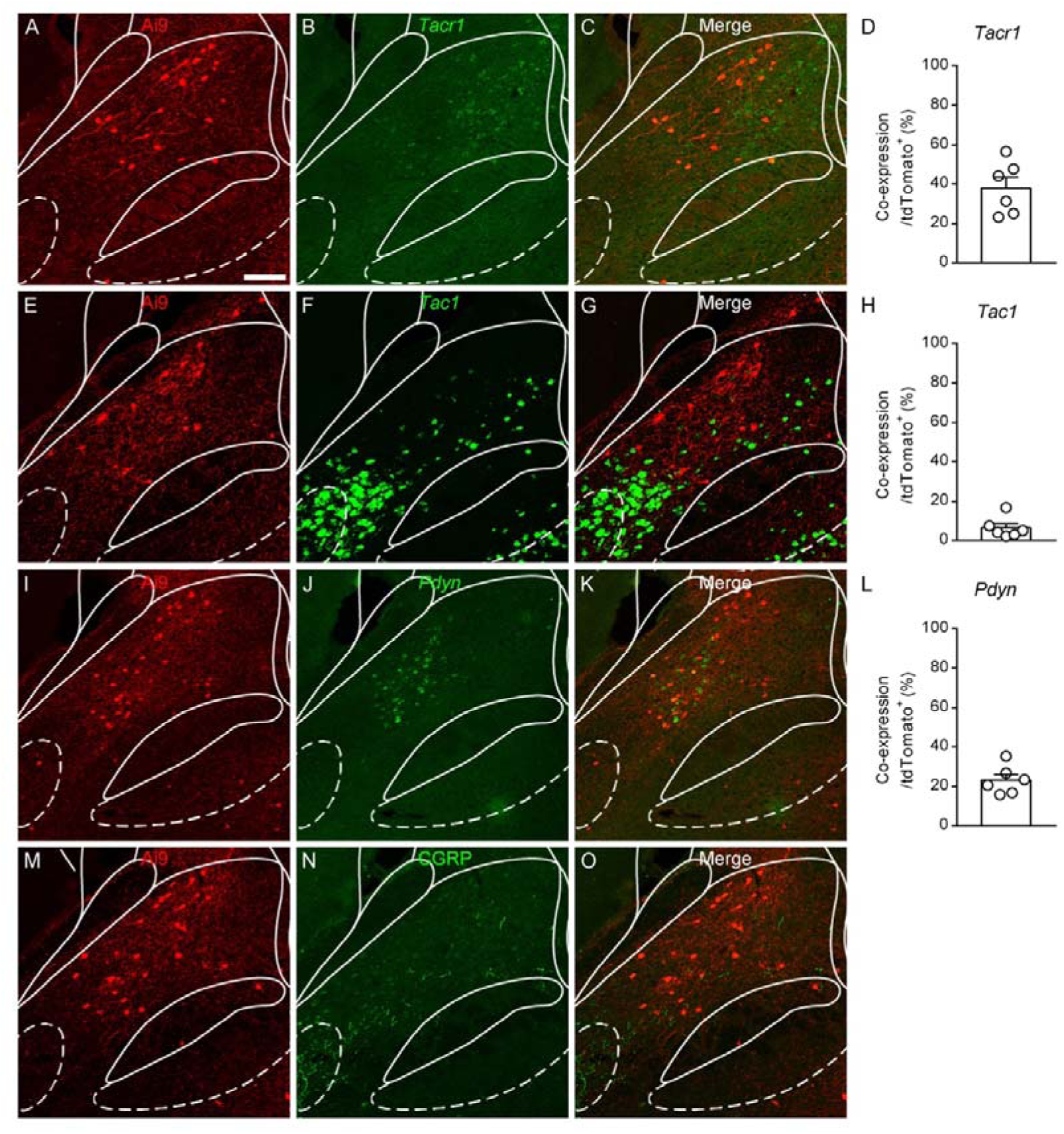
Characterization of PVT-projecting neurons in the PBN nucleus. (A−O) Double staining of tdTomato signals (red) with *Tacr1* mRNA (A−C), *Tac1* mRNA (E−G), *Pdyn* mRNA (I−K), and CGRP protein (M−O). Scale bar: 100 μm. (D, H and L) The proportions of co-expressing neurons of tdTomato positive neurons, *n* = 6 sections from 3 mice. Tacr1, tachykinin 1 receptor; Tac1, tachykinin 1; Pdyn, prodynorphin; CGRP, calcitonin gene-related peptide.

**Figure 1−figure supplement 2.**
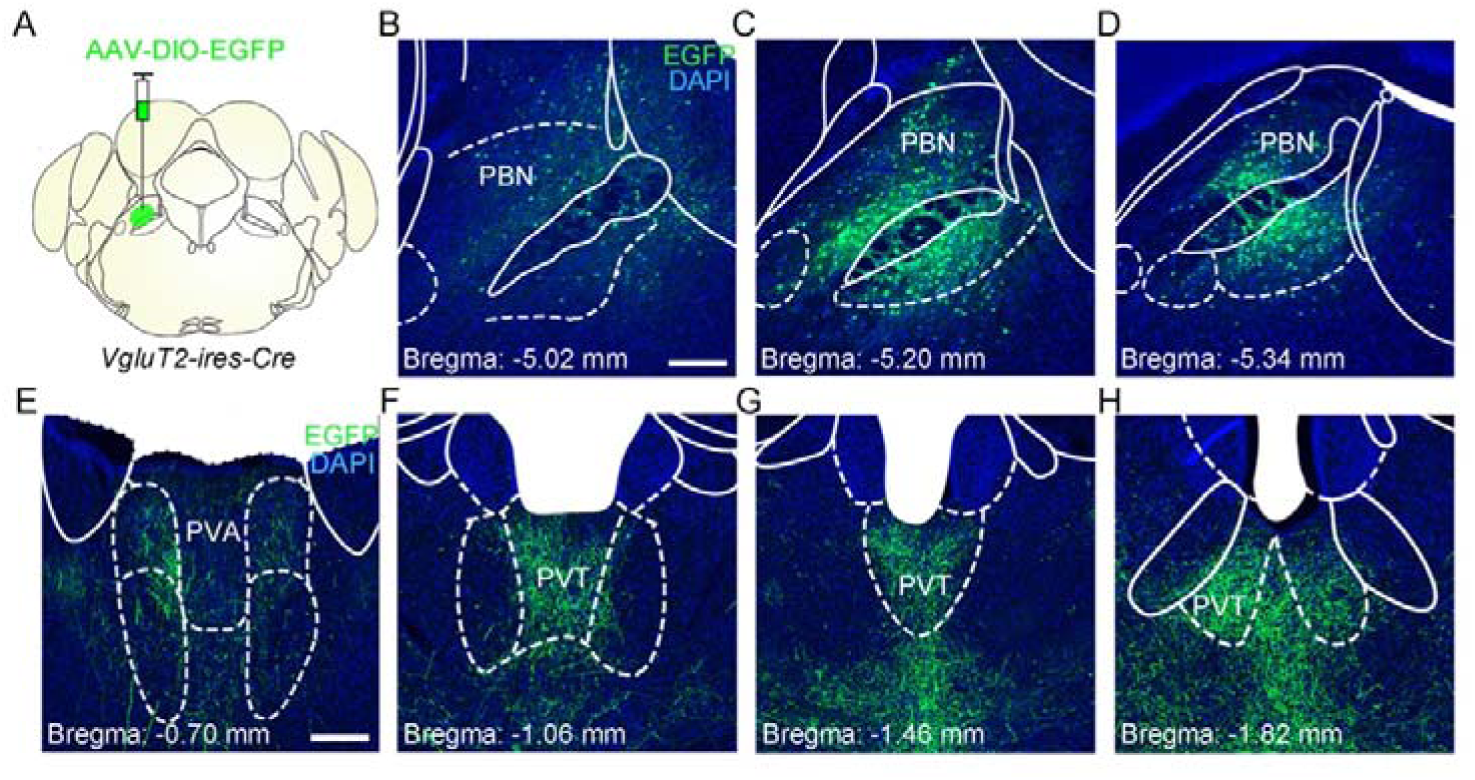
The distribution pattern of the PBN-PVT glutamatergic projection. (A) The illustration for virus injection of AAV2/8-EF1a-DIO-EGFP into the PBN nucleus on *VgluT2-ires-Cre* mice. (B−D) The virus expression in the anterior (B), the middle (C), and the posterior PBN (D). Scale bar: 200 μm. (E−H) The distribution pattern of PBN glutamatergic projection fibers in the anterior (E and F), the middle (G), and the posterior PVT (H). PVA, anterior paraventricular thalamus. Scale bar: 200 μm.

**Figure 1−figure supplement 3.**
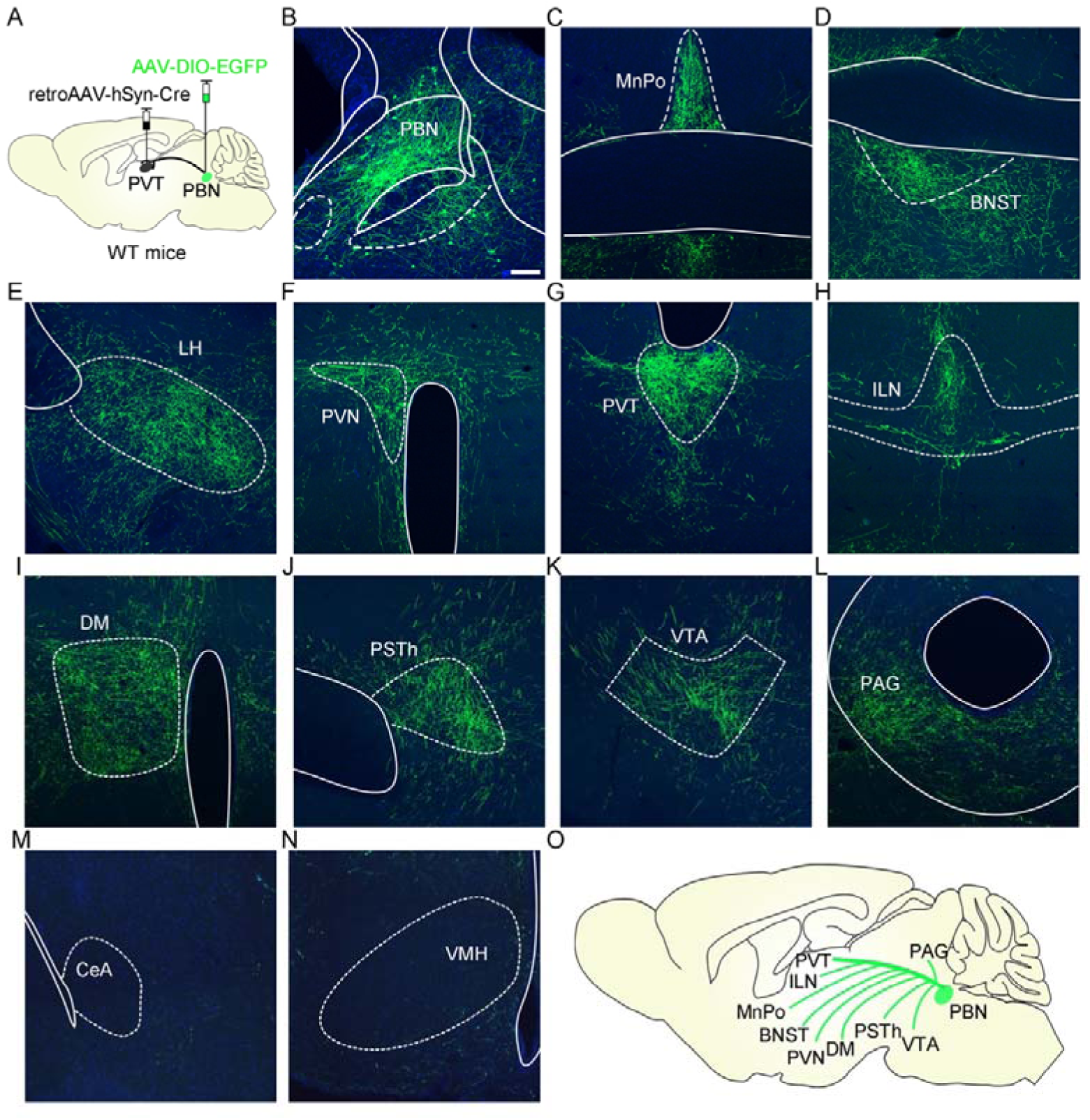
The distribution pattern of collateral projection fibers from PVT-projecting PBN neurons. (A) The illustration showed the injection of retroAAV2/2-hSyn-Cre into the PVT and AAV2/8-EF1a-DIO-EGFP into the PBN to label the PVT-projecting PBN neurons. (B) Examples of AAV2/8-EF1a-DIO-EGFP expression in the PBN. Scale bar: 200 μm. (C−N) The efferents from the PVT-projecting PBN neurons could be found in the MnPo (C), BNST (D), LH (E), PVN (F), PVT (G), ILN (H), DM (I), PSTh (J), VTA (K) and PAG (L) but not in the CeA (M) and VMH (N). (O) Schematic showing summary of the distribution pattern of fibers from PVT-projecting PBN neurons. MnPo, Median preoptic nucleus; BNST, bed nucleus of the stria terminalis; LH, lateral hypothalamic area; PVN, paraventricular nucleus of the hypothalamus; ILN, intralaminar thalamic nucleus; DM, dorsomedial hypothalamic nucleus; PSTh, parasubthalamic nucleus; VTA, ventral tegmental areas; PAG, periaqueductal gray; CeA, central nucleus of the amygdala; VMH, ventromedial hypothalamic nucleus.

**Figure 2−figure supplement 1.**
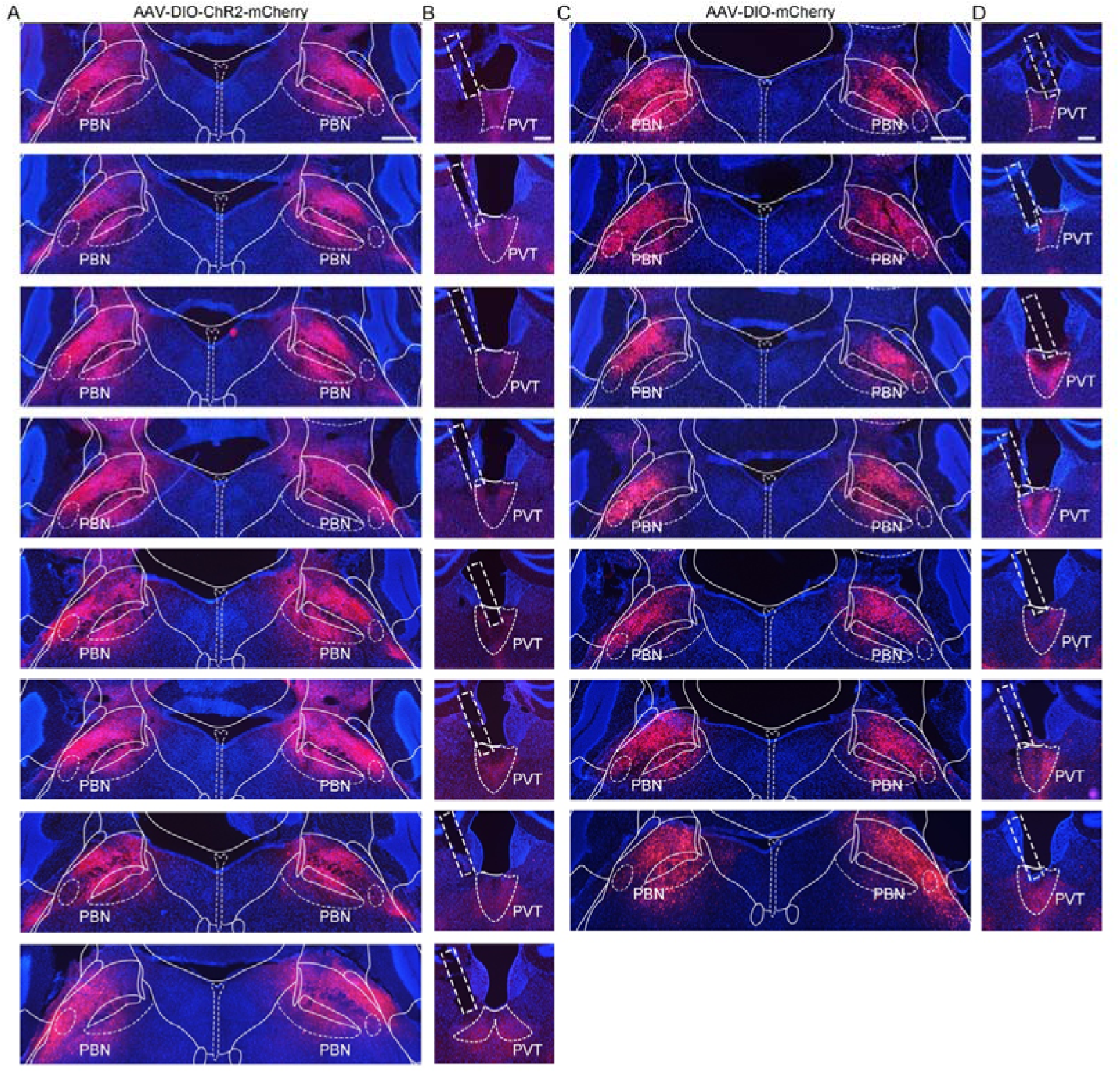
The virus expression in the PBN and the optic fiber position in the PVT of *VgluT2-ires-Cre* mice injected with AAV2/9-EF1a-DIO-ChR2-mCherry virus or AAV2/9-EF1a-DIO-mCherry virus. (A) Histological map showing area of ChR2 expression in the PBN at bregma −5.20 mm in 8 mice. Scale bar: 400 μm. (B) The position of optic fiber (rectangle) in the PVT in the AAV2/9-EF1a-DIO-ChR2-mCherry injected mice. Scale bar: 200 μm. (C) The area of AAV2/9-EF1a-DIO-mCherry virus expression in the PBN at bregma −5.20 mm in 7 mice. Scale bar: 400 μm. (D) Position of the optic fiber tip from 7 mice injected with AAV2/9-EF1a-DIO-mCherry. Scale bar: 200 μm.

**Figure 2−figure supplement 2.**
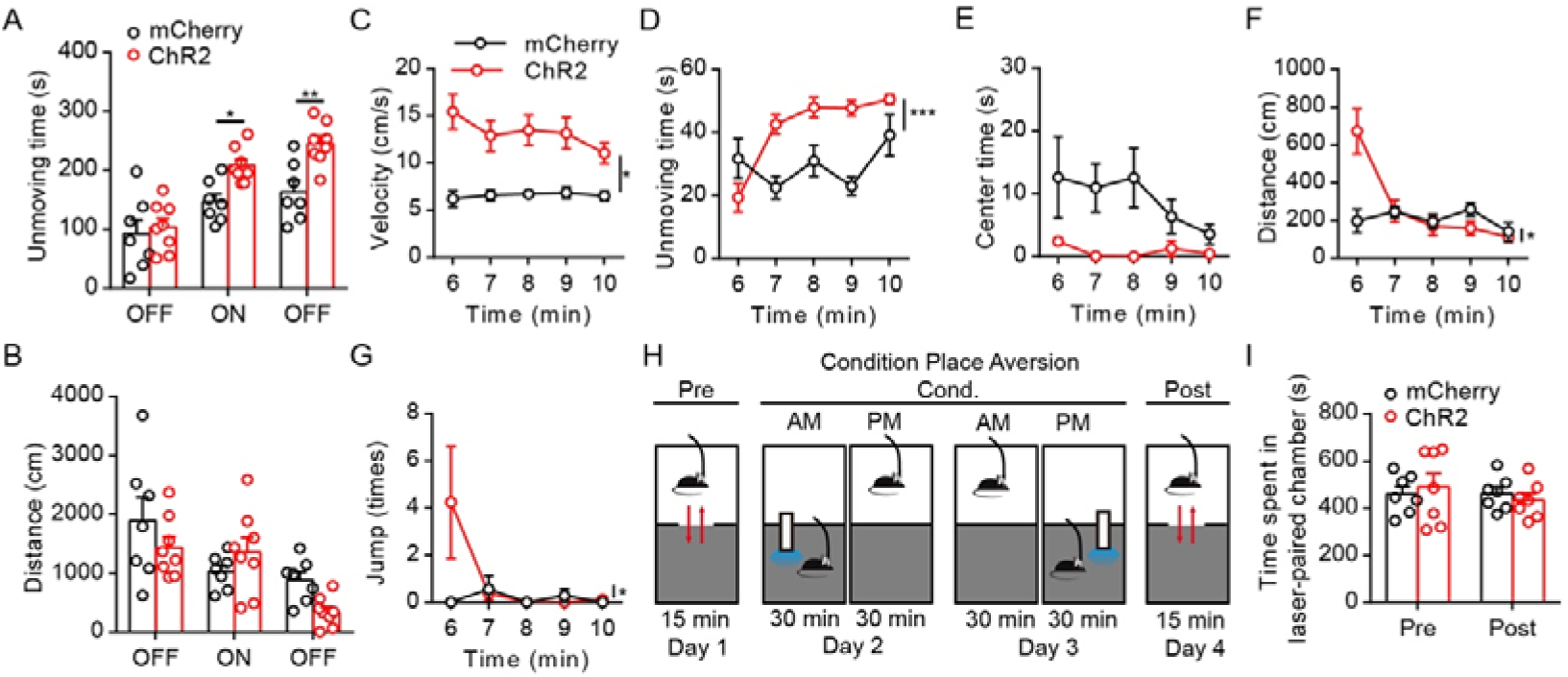
Effects of optogenetic activation of PBN-PVT projection fibers in the OFT and the CPA. (A and B) Quantification of the unmoving time (A) and the distance (B) in the OFT (mCherry group: *n* = 7 mice; ChR2 group: *n* = 8 mice). (C−F) Quantification of the velocity (C), the unmoving time (D), the center time (E), the distance (F) and the number of jumps (G) during the 5−10 minutes laser ON period in the OFT test (mCherry group: *n* = 7 mice; ChR2 group: *n* = 8 mice). (H) Protocol for the prolonged conditioned place aversion (CPA). (I) Photostimulation of PBN-PVT projection did not induce CPA (*n* = 7 mice per group). *p < 0.05, **p < 0.01, ***p < 0.001, all data were represented as mean ± SEM. Two-way ANOVA followed by Bonferroni test for A, B, C, D, E, F, G, and I.

**Figure 3−figure supplement 1.**
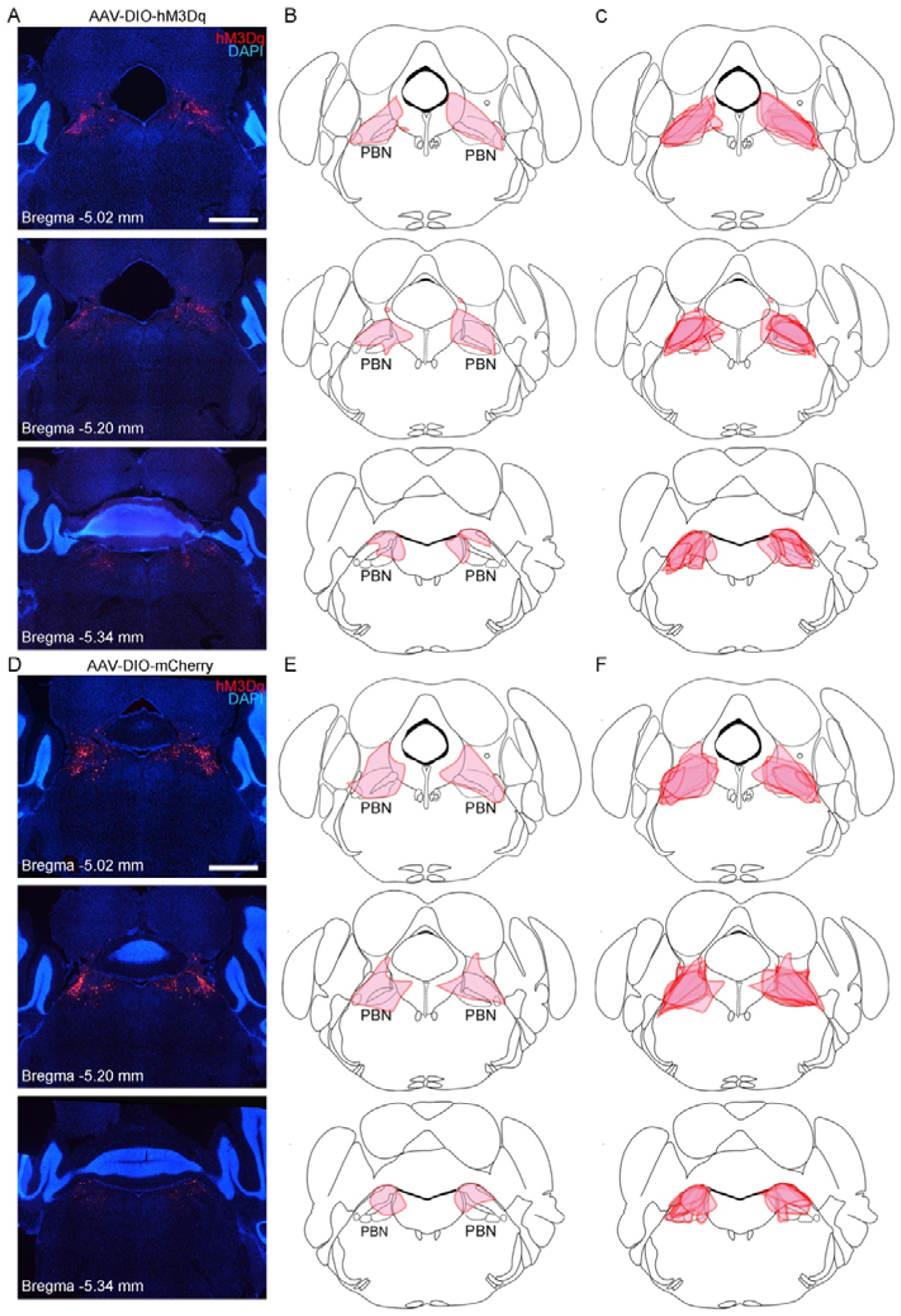
The virus expression in the PBN of mice injected with AAV2/9-hSyn-DIO-hM3Dq-mCherry or AAV2/9-EF1a-DIO-mCherry in the pharmacogenetic manipulation. (A) Representative histological images of hM3Dq expression in an AAV2/9-hSyn-DIO-hM3Dq-mCherry injected mouse at brain level from bregma −5.02 mm to bregma −5.34 mm. Scale bar: 1 mm. (B) Depiction of virus infection area according to the histological images in (A). (C) Superimposed depiction of virus transduction from 8 mice. (D) Representative histological images of mCherry expression in an AAV2/9-EF1a-DIO-mCherry injected mice at brain level from bregma −5.02 mm to bregma −5.34 mm. Scale bar: 1 mm. (E) Depiction of virus infection area according to the histological images in (D). (F) Superimposed depiction of virus transduction from 8 mice.

**Figure 3−figure supplement 2.**
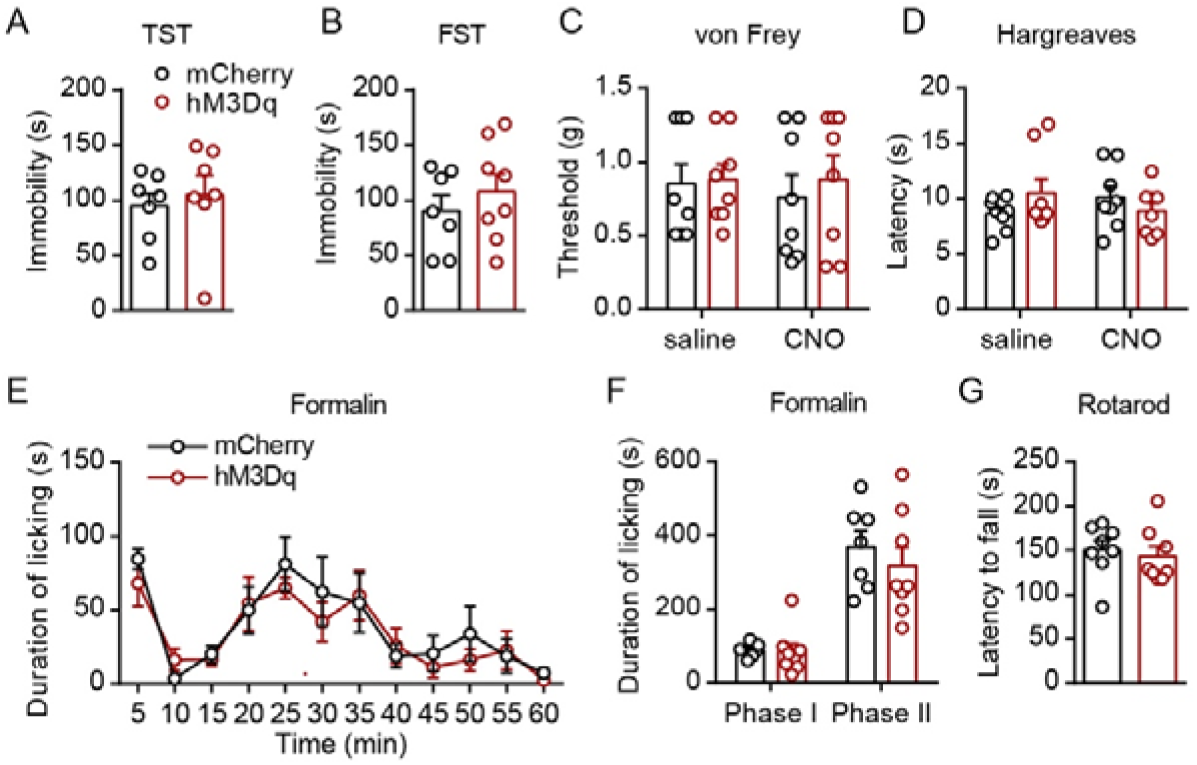
Pharmacogenetic activation of PVT-projecting PBN neurons did not affect depressive-like behaviors, basal nociceptive thresholds, formalin-induced licking behavior, or motor function. (A) Immobility time in the tail suspension test (TST), *n* = 7 mice per group. (B) Immobility time in the forced swimming test (FST), mCherry group: *n* = 7 mice; hM3Dq group: *n* = 8 mice. (C and D) Effects of pharmacogenetic activation of PVT-projecting PBN neurons on the nociceptive response tested by von Frey (C) and Hargreaves (D), *n* = 8 mice per group. (E and F) Duration of licking behaviors in the formalin-induced inflammatory pain test, mCherry group: *n* = 7 mice; hM3Dq group: *n* = 8 mice. Phase I: 0−10 minutes, Phase II: 10−60 minutes. (G) The latency to fall in the rotarod test, *n* = 8 mice per group. Data were represented as mean ± SEM. Unpaired student’s *t*-test for A, B and G. Two-way ANOVA followed by Bonferroni test for C, D, E, and F.

**Figure 4−figure supplement 1.**
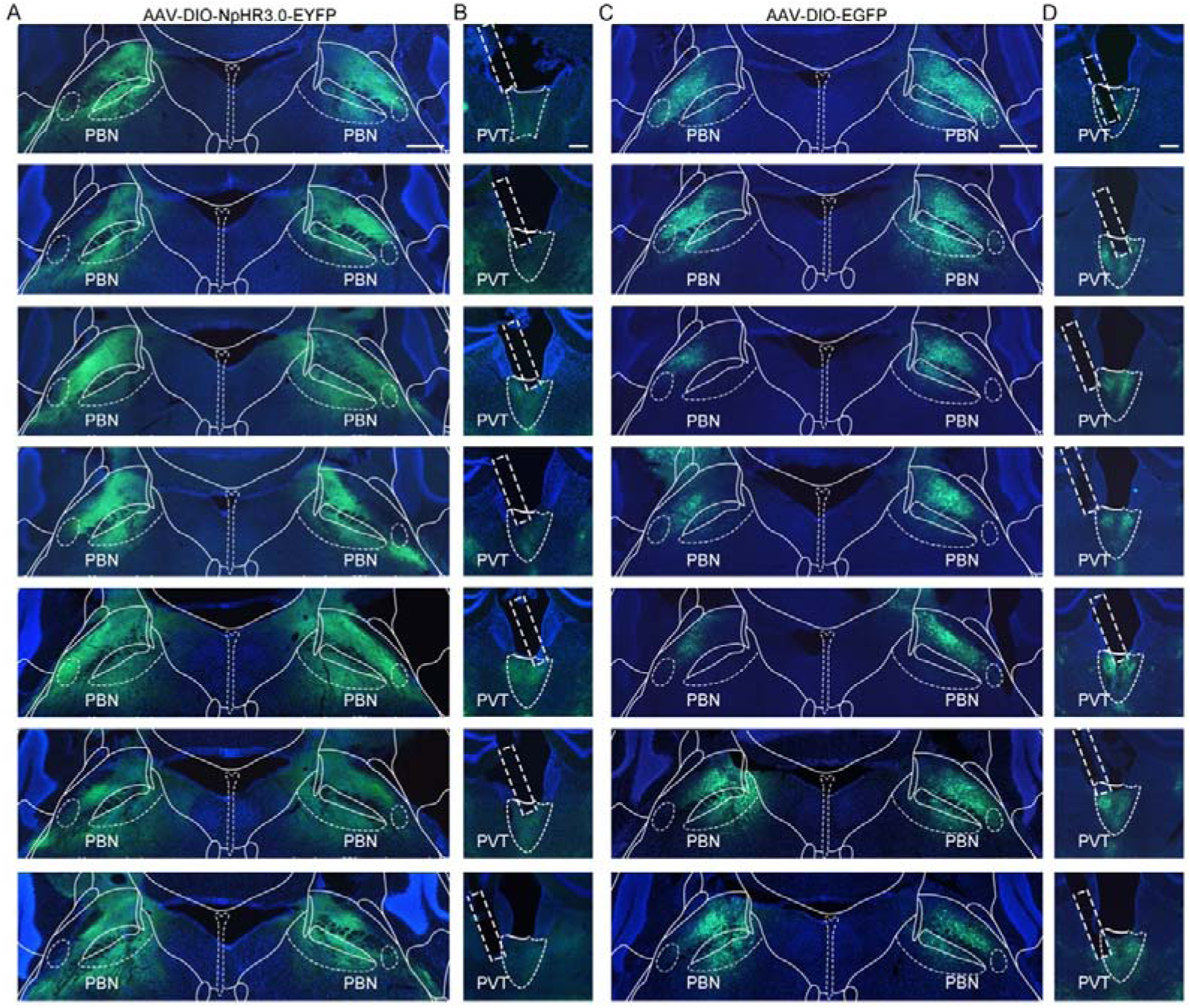
The virus expression in the PBN and the optic fiber position in the PVT of *VgluT2-ires-Cre* mice injected with AAV2/9-EF1a-DIO-NpHR3.0-EYFP or AAV2/8-EF1a-DIO-EGFP. (A) Histological map showing area of NpHR3.0 expression in the PBN at bregma −5.20 mm in 7 mice. Scale bar: 400 μm. (B) The position of optic fiber (rectangle) in the PVT in the AAV2/9-EF1a-DIO-NpHR3.0-EYFP injected mice. Scale bar: 200 μm. (C) The area of AAV2/8-EF1a-DIO-EGFP expression in the PBN at bregma −5.20 mm in 7 mice. Scale bar: 400 μm. (D) Position of the optic fiber tip from 7 mice injected with AAV2/9-EF1a-DIO-mCherry. Scale bar: 200 μm.

**Figure 4−figure supplement 2.**
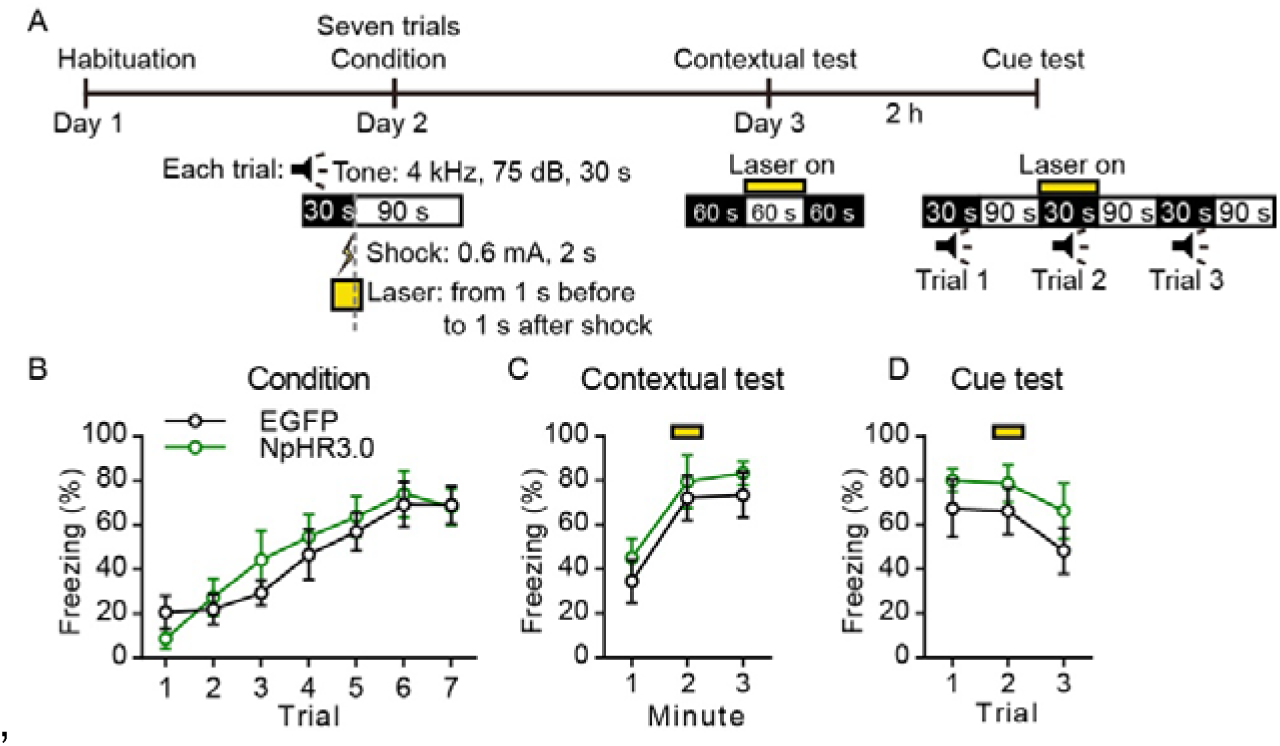
Optogenetic inhibition of the PBN-PVT projection did not affect associative fear memory acquisition and retrieval. (A) The protocol of auditory fear conditioning experiments with optogenetic inhibition of the PBN-PVT projection. (B−D) Quantification of freezing levels during condition trials (B), contextual test (C), and cue test (D), *n* = 7 mice per group. The yellow box indicated optogenetic inhibition. All data were represented as mean ± SEM, two-way ANOVA followed by Bonferroni test for B, C and D.

**Figure 4−figure supplement 3.**
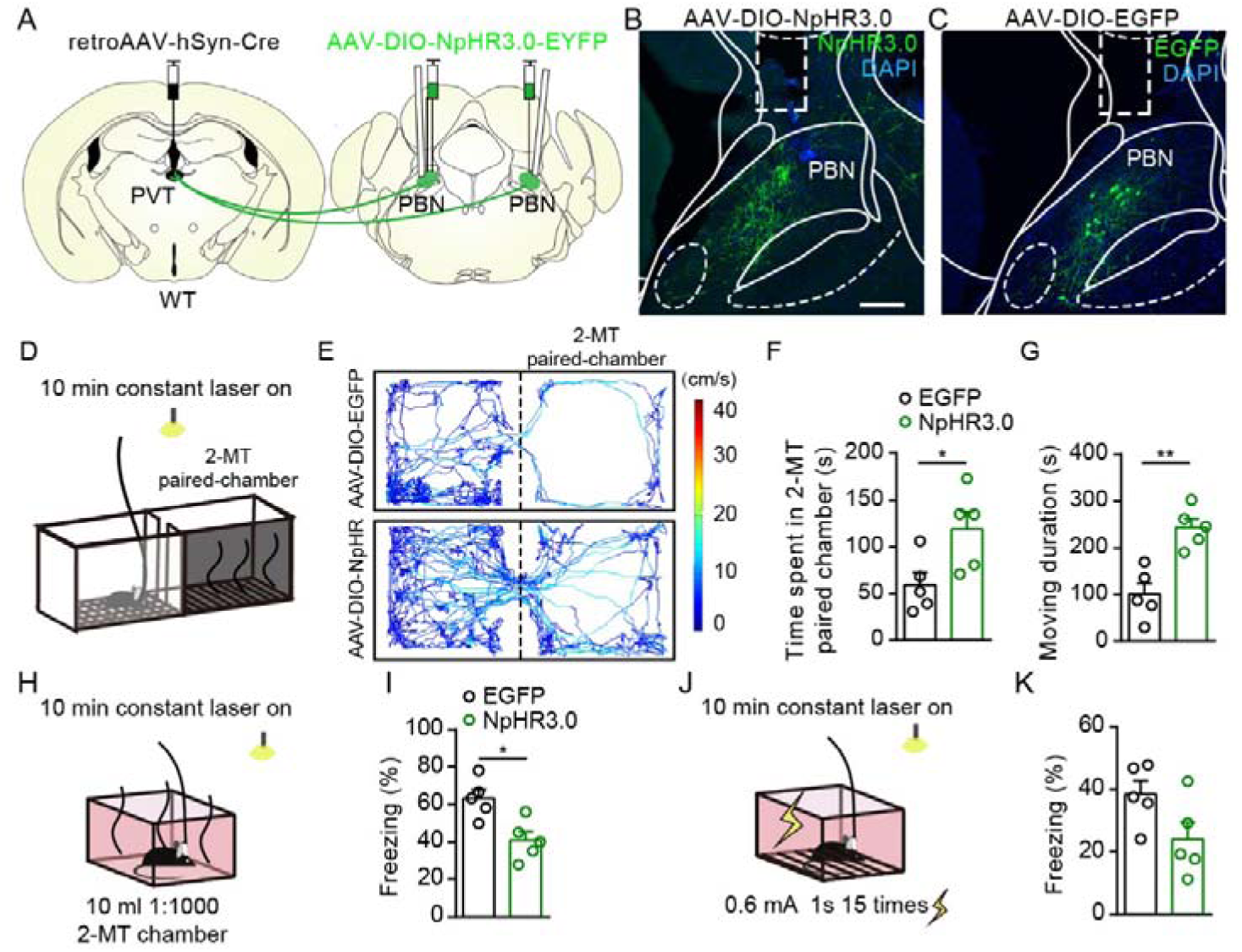
Optogenetic inhibition of the PVT-projecting PBN neurons reduced the aversion-like behavior and fear-like freezing behavior. (A) The illustration showed virus injection of retroAAV2/2-hSyn-Cre into the PVT, bilateral injection of AAV2/9-EF1a-DIO-NpHR3.0-EYFP into the PBN, and bilateral placement of optic fiber above the PBN on WT mice. (B and C) Examples of AAV2/9-EF1a-DIO-NpHR3.0-EYFP (B) and AAV2/8-EF1a-DIO-EGFP (C) expression in the PBN, the rectangle represented the position of the optic fiber. Scale bar: 200 μm. (D) Schematic of 2-MT induced aversion test with optogenetic inhibition via a 589 nm laser. (E) Representative traces of the mice infected with AAV2/8-EF1a-DIO-EGFP or AAV2/9-EF1a-DIO-NpHR3.0-EYFP in the chamber. (F and G) Quantification of the time spent in the 2-MT paired chamber (F) and the moving duration (G), *n* = 5 mice per group. (H) Schematic of 2-MT induced fear-like freezing behavior with optogenetic inhibition via a 589 nm laser. (I) Quantification of the freezing behavior, *n* = 5 mice per group. (J) Illustration of footshock-induced freezing behavior with optogenetic inhibition via a 589 nm laser. (K) Quantification of the freezing behavior, *n* = 5 mice per group.*p < 0.05, **p < 0.01, all data were represented as mean ± SEM. Unpaired student’s *t*-test for F, G, I, and K.

**Figure 5−figure supplement 1.**
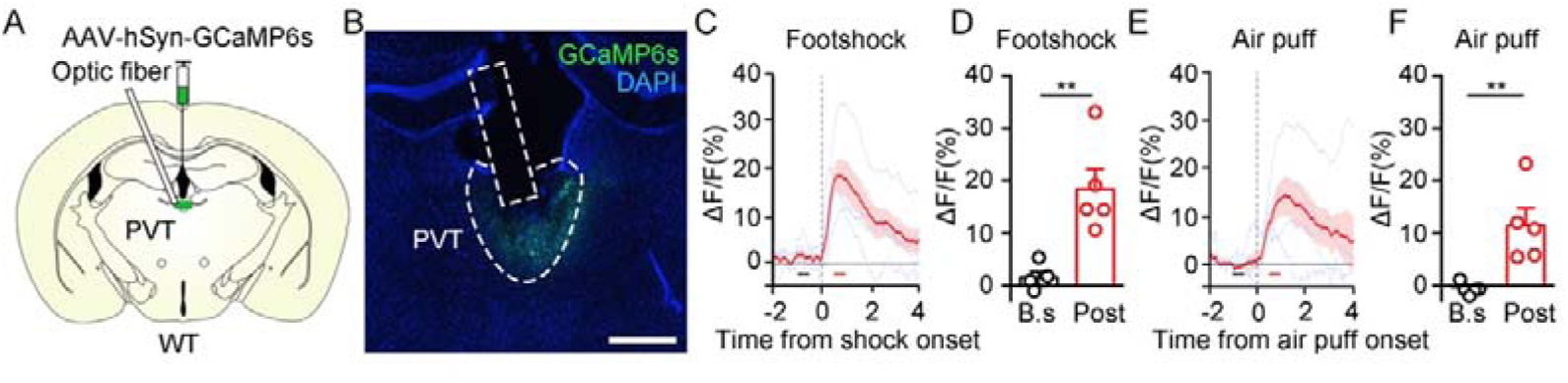
Calcium signals of PVT neurons in response to aversive stimuli. (A) Schematic showed injection of AAV2/8-hSyn-GCaMP6s into the PVT and placement of the optic fiber above the PVT. (B) Representative of GCaMP6s expression and the position of optic fiber in the PVT. Scale bar: 400 μm. (C and D) The calcium signal of the PVT neurons (C) and the quantification of the average Ca^2+^ signal before and after footshock (D). The black bar represented the baseline period (B.s, -1 to -0.5 s), and the red bar represented the post-stimulus period (Post, 0.5 to 1 s), *n* = 5 mice. (E and F) The calcium signal of the PVT neurons (E) and the quantification of average Ca^2+^ signal before and after air puff (F), *n* = 5 mice. **p < 0.01, all data were represented as mean ± SEM. Paired student’s *t*-test for D and F.

**Figure 6−figure supplement 1.**
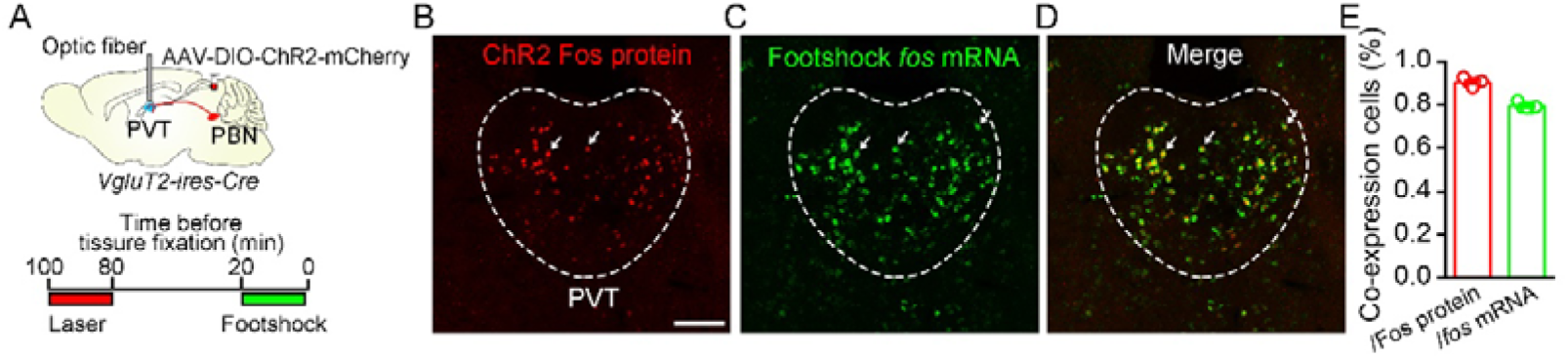
Dual Fos staining detecting Fos protein and *fos* mRNA induced by laser stimulation and footshock. (A) Top: Schematic showed injection of AAV2/9-EF1a-DIO-ChR2-mCherry into the PBN and placement of the optic fiber above the PVT of *VgluT2-ires-Cre* mice. Bottom: Time windows containing laser (20 Hz, 5 mW, 5ms) and shock stimuli (0.5 mA, 1 s, 30 times), separated by 60 minutes of the rest period. (B−D) Example of Fos protein and *fos* mRNA expression in the PVT. Red fluorescence represents Fos protein induced by laser stimulus, and green fluorescence indicates *fos* mRNA detected by *in situ* hybridizations. Scale bar: 100 μm. (E) The proportion of co-expression neurons over Fos protein-expressing cells and *fos* mRNA-expressing cells, *n* = 5 mice.

**Figure 7−figure supplement 1.**
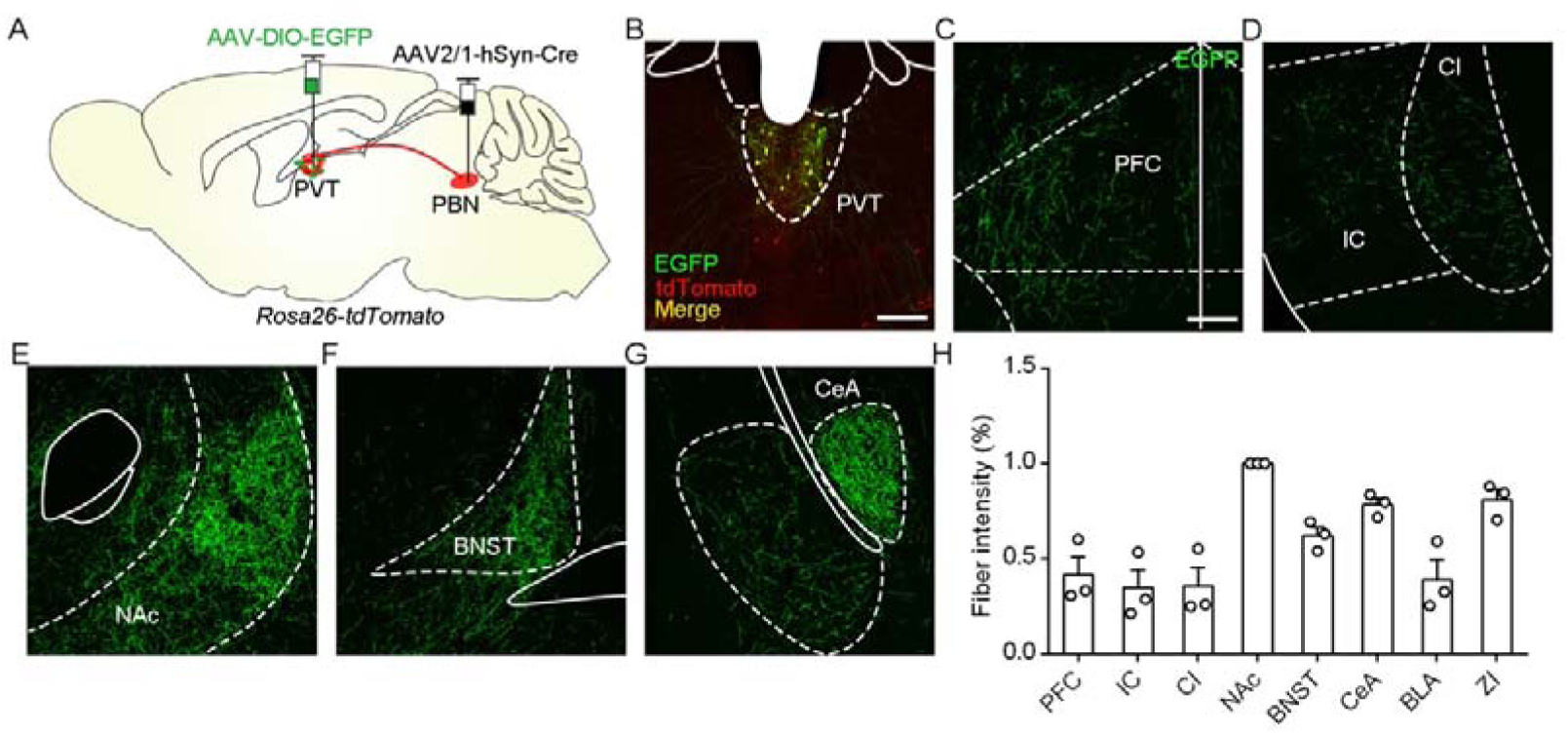
Distribution pattern of projection fibers of PVT_PBN_ neurons. (A) The illustration showed injection of AAV2/1-hSyn-Cre into the PBN and AAV2/8-EF1a-DIO-EGFP into the PVT of *Rosa26-tdTomato* mice. (B) The representative image of EGFP and tdTomato-transduced neurons in the PVT. Scale bar: 200 μm. (C−G) Distribution patterns of EGFP fibers in the PFC (C), CI (D), NAc (E), BNST (F), and CeA (G). PFC, prefrontal cortex; IC, insular cortex; CI, claustrum; NAc, nucleus accumbens core; BNST, bed nucleus of the stria terminalis; CeA, central nucleus of the amygdala. Scale bar: 200 μm. (H) Quantification of the fiber intensity in these brain regions, *n* = 3 mice.

